# Immunogenicity and Protective Efficacy of an Intranasal Live-attenuated Vaccine Against SARS-CoV-2 in Preclinical Animal Models

**DOI:** 10.1101/2021.01.08.425974

**Authors:** Jun-Guy Park, Fatai S. Oladunni, Mohammed A. Rohaim, Jayde Whittingham-Dowd, James Tollitt, Bakri M Assas, Wafaa Alhazmi, Abdullah Almilaibary, Munir Iqbal, Pengxiang Chang, Renee Escalona, Vinay Shivanna, Jordi B. Torrelles, John J Worthington, Lucy H. Jackson-Jones, Luis Martinez-Sobrido, Muhammad Munir

## Abstract

The global deployment of an effective and safe vaccine is currently a public health priority to curtail the coronavirus disease 2019 (COVID-19) pandemic caused by severe acute respiratory syndrome coronavirus 2 (SARS-CoV-2). Here, we evaluated a Newcastle disease virus (NDV)-based intranasal vectored-vaccine in mice and hamsters for its immunogenicity, safety and protective efficacy in challenge studies with SARS-CoV-2. The recombinant (r)NDV-S vaccine expressing spike (S) protein of SARS-CoV-2 administrated via intranasal route in mice induced high levels of SARS-CoV-2-specific neutralizing immunoglobulin A (IgA) and IgG2a antibodies and T cell-mediated immunity. Hamsters vaccinated with two doses of vaccine showed complete protection from clinical disease including lung infection, inflammation, and pathological lesions after SARS-CoV-2 challenge. Importantly, a single or double dose of intranasal rNDV-S vaccine completely blocked SARS-CoV-2 shedding in nasal turbinate and lungs within 4 days of vaccine administration in hamsters. Taken together, intranasal administration of rNDV-S has the potential to control infection at the site of inoculation, which should prevent both the clinical disease and transmission to halt the spread of the COVID-19 pandemic.

## INTRODUCTION

Coronavirus disease 2019 (COVID-19), caused by the severe acute respiratory syndrome coronavirus 2 (SARS-CoV-2), emerged in a seafood market in Wuhan, China during late 2019 and was linked to pneumonia-associated illness, which can culminate in respiratory failure^1^. COVID-19 can be also manifested by pathologies in the cardiovascular and gastrointestinal system and induce a hyperinflammatory syndrome^2-4^. People with immunosuppression, older age (>60s), and with existing co-morbidities such as obesity, diabetes, and hypertension, among others, are highly vulnerable and may bear severe COVID-19 clinical symptoms^5^. COVID-19 has currently overwhelmed healthcare systems around the world with >75 million infections and hundreds of thousands of deaths with a case-fatality rate of approximately 4%^7^. The enormous impact of COVID-19 on lives, and its socioeconomic effect on the human population requires the development of a vaccine that not only reduces the severity of the SARS-CoV-2 infection, but also curbs virus transmission and protects vaccinated people from infection.

SARS-CoV-2 is a positive sense, single-stranded RNA virus with a genome of approximately 30,000 nucleotides. Most of the genome (2/3) encodes for non-structural proteins (nsp) which play critical roles in viral replication and RNA synthesis^7^. The rest of the one-third viral genome encodes the main structural proteins including spike (S), envelope (E), membrane (M), and nucleocapsid (N) proteins. The S protein constitute a 180 kDa homo-trimeric class I viral fusion protein on the surface of virions, which interacts with the cellular carboxypeptidase angiotensin-converting enzyme 2 (ACE2) facilitating SARS-CoV-2 infection of host cells. Owing to S-ACE2 interaction as a solo entry mechanism for SARS-CoV-2, and that the S protein contains several crucial B- and T-cell epitopes^8,9^, it remains the main target for neutralizing antibodies (NAbs) and thus, it serves as a promising vaccine target against coronaviruses, including SARS-CoV-2.

Since the release of SARS-CoV-2 genome sequence, a range of vaccine platforms were repurposed principally targeting the S protein of SARS-CoV-2^10^. Currently, a number of candidate SARS-CoV-2 vaccines are either under development including viral vectored-, inactivated virion- and recombinant protein-based vaccines or being authorised for only emergency use such as lipid nanoparticle encapsulated mRNA vaccine^11,12^. While each of these vaccine platforms present their merit and demerits, vectored-based vaccines possess an enormous potential at eliciting strong and long-lasting adaptive immunity including antibody (Ab) responses^11,12^. Most of the currently developed SARS-CoV-2 vectored-based vaccines adapt the intramuscular route of administration; however, it has been demonstrated that the intranasal route offers superior induction of local mucosal immunity and protection against SARS-CoV-2^13^.

Indeed, owing to the nature of the virus biology, the Newcastle disease virus (NDV) vector offers multifaceted and superior advantages for intranasal immunization. As an avian virus, NDV is fully attenuated in human and non-human primates and causes no clinical symptoms except occasional flu-like short-lived conjunctivitis in humans^14^. Several strains of NDV have been proven to be safe, being used extensively as oncolytic agents in humans^14^. NDV is genetically stable and can tolerate insertion of foreign transgenes of approximately 5 Kb into its genome without compromising viral replication^14^. Noticeably, humans typically lack pre-existing immunity against NDV, which favours NDV over currently applied human viral vectors such as human adenovirus (Ad), measles virus (MeV), modified vaccinia Ankara (MVA) or vesicular stomatitis virus (VSV). Owing to its tendency to replicate in the respiratory tract, NDV has also been shown to induce local mucosal immunity offering an additional advantage of interfering with viral shedding. Importantly, NDV-based vaccines can be produced in chicken embryonated eggs or in US Food and Drug Administration (FDA)-approved cell lines using protocols and infrastructure currently being applied for influenza virus vaccines. Given these and other features, apathogenic strains of NDV have been used as live attenuated vaccines (LAVs) against multiple viruses including influenza^15,16^, SARS-CoV^17^, human immunodeficiency virus^18,19^, human parainfluenza^20,21^, rabies^22^, Nipah^23^, Rift Valley fever^24^ and Ebola^25-27^. Notably, NDV-based vaccines have been shown to be safe and effective in multiple animal models including mice^16^, dogs^22^, pigs^23^, cattle^22^, sheep^28^, African green and rhesus monkeys^15,17^ and ultimately humans^17, –29-31^.

We have previously developed an NDV-based SARS-CoV-2 vaccine encoding a human codon optimized full length spike (S) protein of SARS-CoV-2 (rNDV-S) using reverse genetics^33^. rNDV-S replicated in chicken embryonated eggs and cultured cells to levels comparable to a recombinant wild-type NDV (rNDV-WT). Here we tested the immunogenicity and safety of this rNDV-S-based LAV in mice and advanced our studies to assess protective efficacy in hamsters. The rNDV-S induced robust systemic humoral and cell-mediated immune responses with two intranasal vaccine doses in mice and fully protected against lung infection, inflammation, and pathology after SARS-CoV-2 challenge in hamsters. Intranasal administration of two doses of rNDV-S induced high levels of SARS-CoV-2 NAbs and anti-SARS-CoV-2 immunoglobulin A (IgA) and IgG2a and conferred complete protection against SARS-CoV-2 infection. Importantly, single or double dose of rNDV-S completely blocked the virus shedding into nasal turbinate and lungs of hamsters. Altogether, intranasal immunization of rNDV-S has the potential to control SARS-CoV-2 infection at the site of inoculation, which should prevent both virus-induced disease and transmission, representing an excellent valid option for the prevention of SARS-CoV-2 infection and associated COVID-19 disease.

## RESULTS

### Intranasal vaccination with rNDV-S successfully establishes a viral load with no accompanying adverse pathology

In order to explore the potential of intranasal live-attenuated vector vaccine against SARS-CoV-2, we engineered rNDV-S (*i*.*e*., avian orthoavulavirus 1, AOaV-1) encoding a human codon-optimized SARS-CoV-2 full-length S glycoprotein gene, including the ectodomain, transmembrane domain, and cytoplasmic domain. The S gene was cloned in a pre-optimized gene junction between phosphoprotein and matrix gene of NDV. As we demonstrated previously^33^, the rNDV-S replicated comparably to rNDV-WT in both cell culture and avian eggs and spreads within cells independently of exogenous trypsin, proposing a competitive vaccine candidate.

For safety and immunogenicity assessment of rNDV-S vaccine in mice, groups of 12-week-old BALB/c mice were immunized by intranasal inoculation with 10^6^ PFU of test vaccine rNDV-S or wild type NDV (rNDV-WT) or were mock-vaccinated with phosphate buffer saline (PBS) (Fig. 1a). The rNDV-WT and one rNDV-S group of mice received a booster dose of 10^6^ PFU of rNDV-WT or rNDV-S, respectively, a week later, while other groups were mock-boosted. All animals were euthanized on day 19 post-vaccination for safety assessment, as well as pathological and immunological responses. Mice were monitored daily for weight loss, health status, and feed intake. Initial starting weights did not significantly differ in any experimental group prior to treatment (Fig. 1b and Supplementary Fig. 1), and there was no significant alteration in the percentage of weight in groups immunised with rNDV-S and the control mock-vaccinated group across the time course of the experiment (Fig.1b). The first dose of rNDV-S induced a significant reduction in percentage of basal weight at 2- and 3-days post-vaccination (DPV), but weights returned to comparable levels to mock immunised mice on the following day and for the rest of the time course (Fig.1b). A reduction in percentage weight was also seen at 14 and 15 DPV in the rNDV-S (prime + boost) group, but weights also quickly returned to mock levels for the rest of the study. Analysis of daily chow intake revealed no significant differences in daily chow consumption among groups (Fig. 1c). Mice were scored daily assessing movement, hair coat, temperature, eyes, and signs of hunching on a 14-point scale. No adverse clinical disease signs were observed with indicated daily score “0” in the duration of experiments. Following the end of the experiment at day 19 post-vaccination, the gross and histopathological assessment of right lung lobe showed no significant lesions in any of the treated groups (Fig. 1d, e). Further to confirm that intranasally delivered rNDV-S replicated in immunised mice, we examined the expression of matrix (M) gene of NDV in various tissues at 19 DPV using qRT-PCR. Results indicate a high number of NDV copies, and a comparable replication of rNDV-S and rNDV-WT was observed in the respiratory tract (nasal turbinate, trachea, and lungs) (Fig. 1f-h). However, comparatively reduced replication was observed in the gut of mice vaccinated with rNDV-S or rNDV-WT (Fig. 1i). Collectively, these data indicate that rNDV-S intranasal vaccination showed no adverse pathology in the tissues examined and the rNDV-S vaccine replicated significantly in the target tissues comparable to the rNDV-WT.

**Fig 1.**
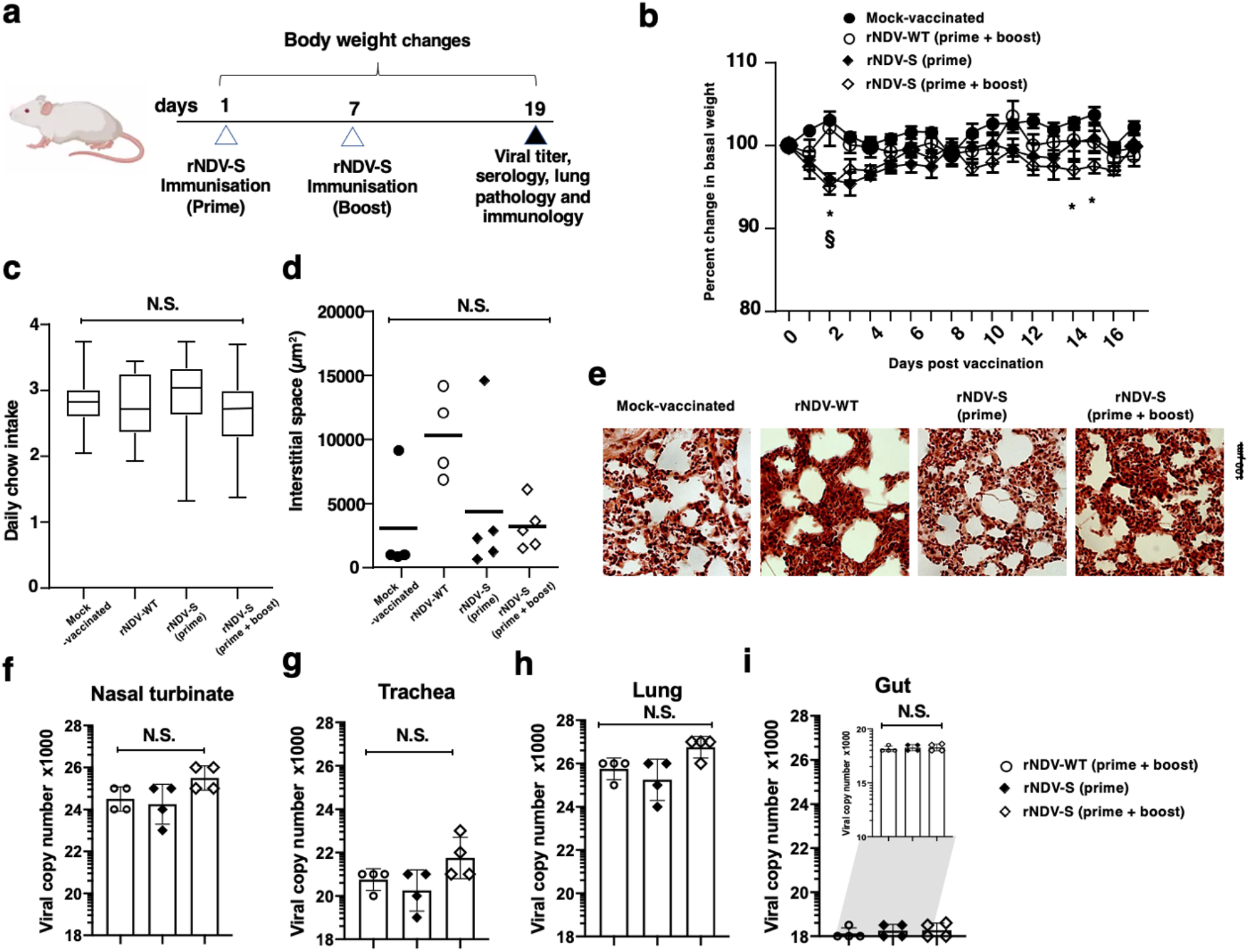
No adverse pathology results following rNDV-S intranasal vaccination. (**a**) Experimental model for the prime and prime + boost intranasal vaccination of mice. (**b**) Percentage weight change of mice following instillation of PBS or indicated rNDV constructs over the experimental time-course. (**c**) Daily chow intake by mock-vaccinated or rNDV-S vaccinated mice. Mean interstitial space size of right lung following H&E staining and assessed via ImageJ (**d**) and representative images (**e**). Viral replication of intranasally administered rNDV in nasal turbinate (**f**), trachea (**g**), lung (**h**) and gut (**i**). Data (n = 4–5 mice/group); *, P<0.05 and NS, non-significant between naïve and vaccinated groups, error bars represent SE of means via repeated t-test or ANOVA with Dunnett’s post-test.

### Intranasal immunization with rNDV-S elicits potent S-protein binding Abs and NAbs in mice

To evaluate the immunogenicity of rNDV-S, sera samples collected at 19 days post-vaccination were used to evaluate the total IgG antibody titers using ELISAs and levels of neutralizing antibodies using microneutralization and pseudo-particle entry inhibition assays, respectively. To detect anti-spike serum Abs, full-length spike (S) protein or recombinant Spike protein receptor binding domain (RBD) expressed using the Drosophila S2 cell system were used to coat ELISA plates. After prime immunization only, the sera from mice immunized with rNDV-S contained S protein-specific Abs, which were significantly increased with the boosting dose. In comparison, no S protein-specific Abs were detected in sera from the mice vaccinated with rNDV-WT or mock-vaccinated (Fig. 2a). Next, we assessed the levels of total IgG responses against purified RBD. The rNDV-S only induced significantly high levels of anti-RBD-specific IgG in the prime + boost vaccinated group when compared to mock-immunised or rNDV-WT-immunised mice (Fig. 2b). We next functionally characterized serum Ab responses by the lentiviral entry inhibition and focal-reduction neutralization tests^34,35^. As expected, serum from mock- or rNDV-WT vaccinated mice did not inhibit the entry of pseudoviral particles (Fig. 2c). In contrast, serum from rNDV-S prime or rNDV-S prime + boost groups significantly inhibited the entry of the pseudoviral particles. Correspondingly, compared to mice mock-immunised or immunised with rNDV-WT, a significant level of neutralization of SARS-CoV-2 infection was observed by sera collected from rNDV-S immunised mice, both prime and prime + boost, the latter more efficient in generating NAbs (Fig. 2d). Overall, rNDV expressing SARS-CoV-2 S protein elicited high titters of protein binding and virus NAbs in mice, especially when a prime + boost regime was utilized.

**Fig 2.**
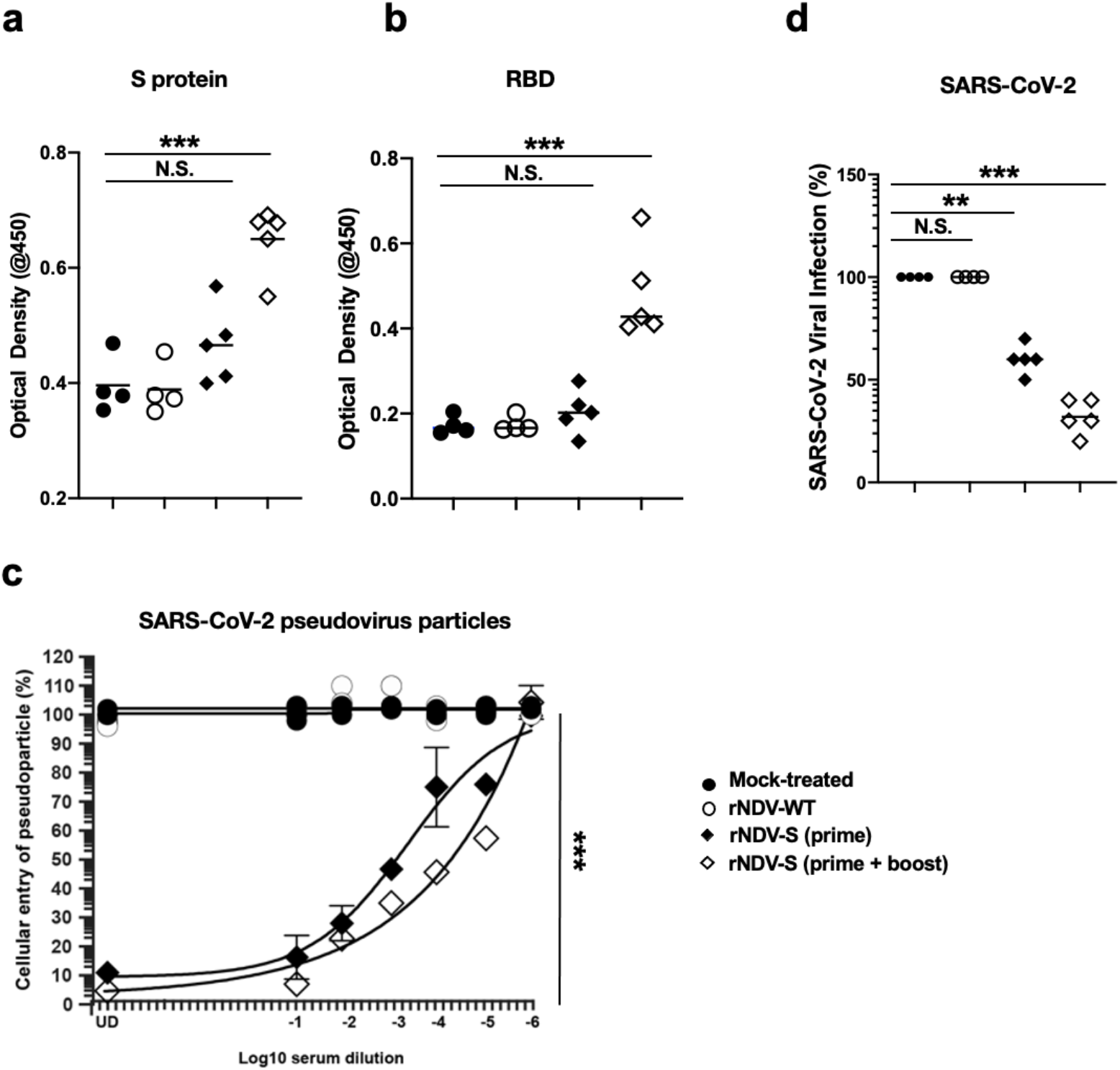
Intranasal administration of rNDV-S elicits high titers of binding and NAbs in mice. (**a**) S-specific serum IgG titers measured by ELISA. Sera from animals at 19 days after-prime and after prime + boost were isolated and used in the ELISA against a recombinant trimeric S protein. Sera from mock-vaccinated mice and mice vaccinated with rNDV-WT were used as control. (**b**) Corresponding sera from all group of mice were used to detect Abs against RBD of spike protein. (**c**) Lentiviruses expressing the S protein of SARS-CoV-2 were used to perform pseudoparticle entry inhibition assay. Sera samples collected from mice after prime or after prime + boost inhibited the entry of lentiviral pseudoparticles compared to sera collected from mice vaccinated with rNDV-WT or mock-vaccinated (**d**) Sera samples from all groups were used to measure neutralization of SARS-CoV-2. Sera from prime from group of mice which were vaccinated with rNDV-S prime only or after prime + boost group of mice showed marked virus neutralization compared to sera from mock-treated or vaccinated with rNDV-WT. Data (n = 4–5 mice/group); *, P<0.05; **, P<0.01; or ***, P<0.005 between naïve and vaccinated groups, error bars represent SE of means via repeated T-test or ANOVA with Dunnett’s.

### Intranasal vaccination with rNDV-S does not induce overt myeloid inflammatory responses in the lungs, pleural cavity or systemically

We next analysed systemic and local immunological responses to determine whether innate and adaptive responses were beyond the basal response induced by rNDV-WT. We did not observe significant alterations in cellularity of the spleen, used as a surrogate marker of systemic inflammation (Supplementary Fig. 2a). We also did not observe significant changes in the rNDV-S prime + boost group percentages of myeloid subsets examined, including neutrophils, monocytes, inflammatory monocytes, or eosinophils (Supplementary Fig. 2b). However, we observed a significant increase in the percentage of dendritic cells (DCs) in the spleen of rNDV-WT immunised animals as compared to the mock vehicle control, and although no significant increase was seen in the rNDV-S prime immunised group, a similar but non-significant trend in DC levels was seen in murine spleen after prime + boost rNDV-S immunisation (Supplementary Fig. 2b). Despite the observed slight morphological changes in the cellular architecture in the lung (Fig. 1d, e), we did observe a significant increase in the cellularity of the lung in the rNDV-S (prime + boost) immunized group, as compared to all other experimental groups studied (Supplementary Fig. 2c). However, when we further examined the myeloid immune subsets, we did not observe significant differences in neutrophils, Ly6C-monocyte/macrophages, alveolar macrophages, and eosinophils as compared to the mock vehicle control group. A significant decrease in the Ly6c+ monocyte/macrophage subset was observed in the rNDV-S (prime + boost) group (Supplementary Fig. 2d). Interestingly and similar to the results in the spleen, we only observed a significant increase in the total DC population in the lung of mice receiving the rNDV-WT vaccine as compared to mock-vaccinated group (Supplementary Fig. 2d).

To investigate further whether a local inflammatory response occurred following intra-nasal vaccination, we assessed the immune response within the pleural cavity. Our results indicate that there were no statistically significant increases in total CD45^+^ haematopoietic cells, neutrophils, eosinophils, monocytes, and F4/80^lo^MHCII^hi^ myeloid cells within either pleural lavage fluid (Supplementary Fig. 3a-f) or pericardial adipose tissue (Supplementary Fig. 3g-k). However, although not significant, a trend for higher amounts of F4/80^hi^MHCII^lo^ pleural cells was observed in rNDV-S prime + boost group (Supplementary Fig. 3f>). These data suggest that there is no lasting abhorrent myeloid inflammatory response generated at the site of vaccination or systemically following intranasal delivery of rNDV-S as compared to the mock-vaccinated mice.

**Fig 3.**
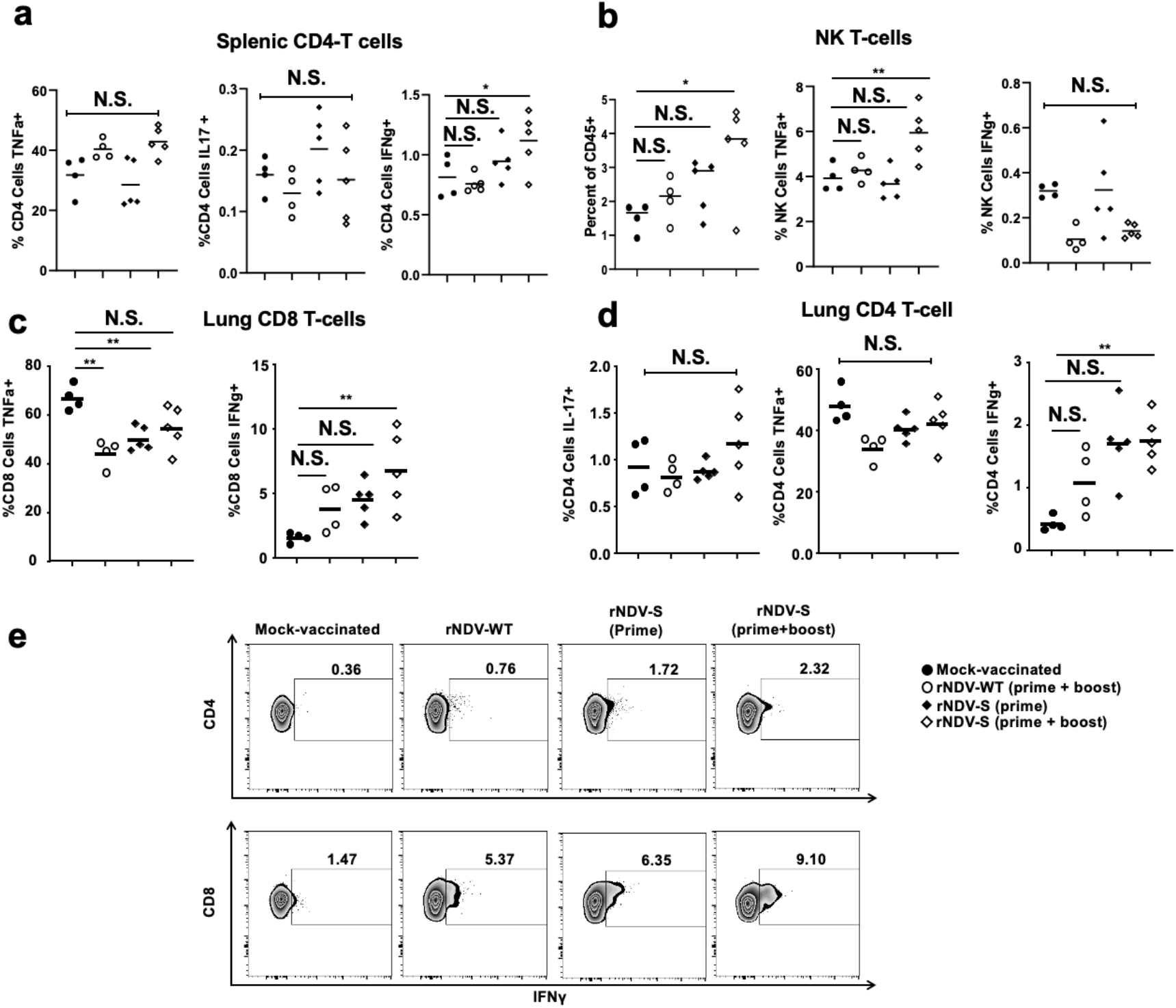
Vaccination with rNDV-S drives both CD4 and CD8 T-cell cytokine responses. Flow cytometric analysis of cytokine responses to SARS-CoV-2 S protein from splenic CD4 (**a**) and NK (**b**) T-cell populations and lung CD4 (**c**) and CD8 (**d**) T cell populations from mice on day 19 following instillation of mock (PBS) or indicated rNDV constructs intranasally on days 0 & 7. (**e**) Representative T-cell flow cytometry plots of lung cytokine production of (**c**) and (**d**). Data (n = 4–5 mice/group); *, P<0.05; **, P<0.01; or ***, P<0.005 between naïve and vaccinated groups, error bars represent SE of means via ANOVA with Dunnett’s post-test.

### Intranasal vaccination with rNDV-S drives T-cell IFNγ responses both in the lung and systemically

We next examined T-cell subsets, asking what antigen specific cytokine responses were produced against the purified full length S protein both systemically and locally at the vaccination site. Systemically, within the spleen, there was no alteration in the percentage of splenic CD4+ or CD8+ T-cell subsets in any of the experimental groups studied (Supplementary Fig. 4a). We also did not see a significant increase in CD8+ T-cell TNF^+^ or CD8+ T-cell IFNγ^+^ cell populations in response to the full-length S protein Ag in any of the groups studied (Supplementary Fig. 4b). Conversely, splenic CD4+ T-cell IFNγ^+^ (Fig. 3a) and NK T-cell TNF^+^ (Fig. 3b) were significantly increased in the rNDV-S boosted group indicating a systemic SARS-CoV-2 S specific response following vaccination with rNDV-S.

**Fig 4.**
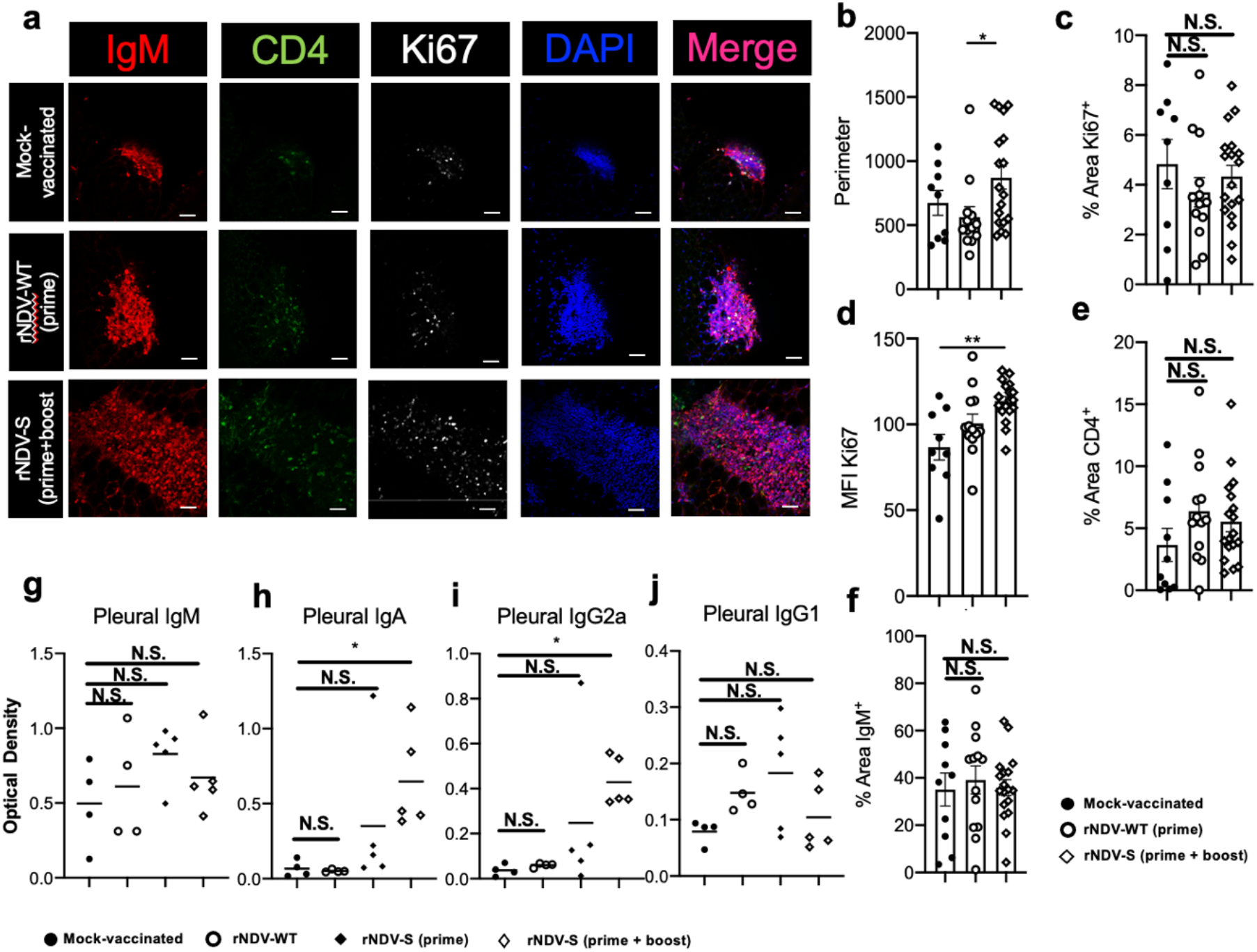
Fat associated lymphoid clusters (FALCs) expand within the pericardium and class switched Abs are present within the pleural fluid following vaccination. Expansion of fat associated lymphoid clusters within the thoracic cavity in response to rNDV-S. (**a**) Whole mount immuno-fluorescence imaging of the murine mediastinum on day 19 following instillation of PBS or indicated rNDV constructs, intranasally, on days 0 & 7. (**b**) Cluster perimeter. (**c)** %Ki67^+^ area of cluster and (**d**) mean fluorescence intensity of Ki67 expression. (**e**) %CD4^+^ area of cluster. (**f**) %IgM^+^ area of cluster. Scale bar = 50μm. IgM (**g**), IgA (**h**), IgG2a (**i**) and IgG1 (**j**) Abs against SARS-CoV-2 spike protein receptor binding domain present within pleural fluid on day 19 following instillation PBS or indicated NDV construct i.n. on day 0 & day 7. Data (n = 4–5 mice/group); *, P<0.05; **, P<0.01; between mock and vaccinated groups, N.S. = non-significant, error bars represent SE of means via one-way ANOVA with Sidak’s post-test.

Locally, within the lung homogenate, a significant decrease in CD4+ T cells and an increase in CD8+ T cell percentage was observed following vaccination with all NDV containing groups (Supplementary Fig. 4c). Looking at cytokine responses we saw a significant reduction in CD8+ T-cell TNFα production in both the NVD-WT and NVD-S groups and a significant increase in IFNγ production in the rNDV-S boosted group as compared to the mock vehicle group (Fig. 3c, e). However, despite an overall decrease of CD4 T cell percentage and no alteration in CD4+ T cell subpopulations producing IL-17 or TNF, a significant increase in CD4+ T cell IFNγ^+^ (6.4-fold) was observed also in the rNDV-S boosted group as compared to the mock-vaccinated group (Fig. 3d, e). Collectively, these data indicate both a systemic and local IFNγ antigen specific T-cell response following intranasal vaccination with rNDV-S.

### Intranasal vaccination with rNDV-S drives NAb responses specifically targeting SARS-COV-2 S protein

Fat associated lymphoid clusters (FALCs) are sites of local Ab production within the pericardium and mediastinum, known for their early response to intranasal challange^36^. Using whole-mount immunofluorescence staining, we were able to detect an expansion of FALCs within the mediastinum at 19 DPV in the rNDV-S boosted group (Fig. 4a, b). Within FALCs of mice receiving rNDV-S boost, and supporting an earlier response, there was a significant increase of the level of Ki67^+^ proliferation, representative of cellular division, (Fig. 4c, d). Although not reaching significance, a trend for increased percentage area of CD4 was seen, suggesting a local accumulation of CD4 T cells within the mediastinum (Fig. 4e). No increase in the % area of IgM (B cell marker) was seen within the pericardium at day 19 DPV (Fig. 4e).

IgM is the first polyclonal Ab induced during an immune response, and following induction of adaptive immunity B cells, IgM class switches to other Ab subclasses defined by the cytokine milieu. As FALCs are the site from which Abs detected in pleural fluid are produced we next confirmed the local activation of B cells by assessing the presence of Abs against SARS-CoV2 S RBD within the pleural fluid. Within the pleural fluid no antigen specific IgM was detected (Fig. 4g), however a significant increase in both RBD-specific IgA (Fig. 4h) and IgG2a (Fig. 4i) but not IgG1 (Fig. 4j) levels were present within the pleural fluid of mice in the rNDV-S boosted group compared to control mice and those receiving rNDV-WT. These data suggest that B cells producing the Abs had class switched and is consistent with the antigen specific T cell derived IFNγ^+^ detected upon re-stimulation, driving a switch to IgA and/or IgG2a (Fig. 4h, i). We next determined the number of B cells present within the pleural lavage fluid and digested pericardium and did not observe significant differences in the numbers of B1a, B1b, or B2 cells present within the pleural lavage fluid or pericardium when comparing rNDV-S boosted to mock control mice (Supplementary Fig. 5). Furthermore, we also did not detect a significant increase in pericardial B cell proliferation (Supplementary Fig. 6a, b), suggesting that Abs detected within the pleural fluid may have been secreted earlier or resulted from an ongoing systemic B cell response within primary lymphoid organs. To further support this, systemic anti-full-length spike and anti-RBD Ab responses were observed which neutralized SARS-CoV-2 (Fig. 2).

**Fig 5.**
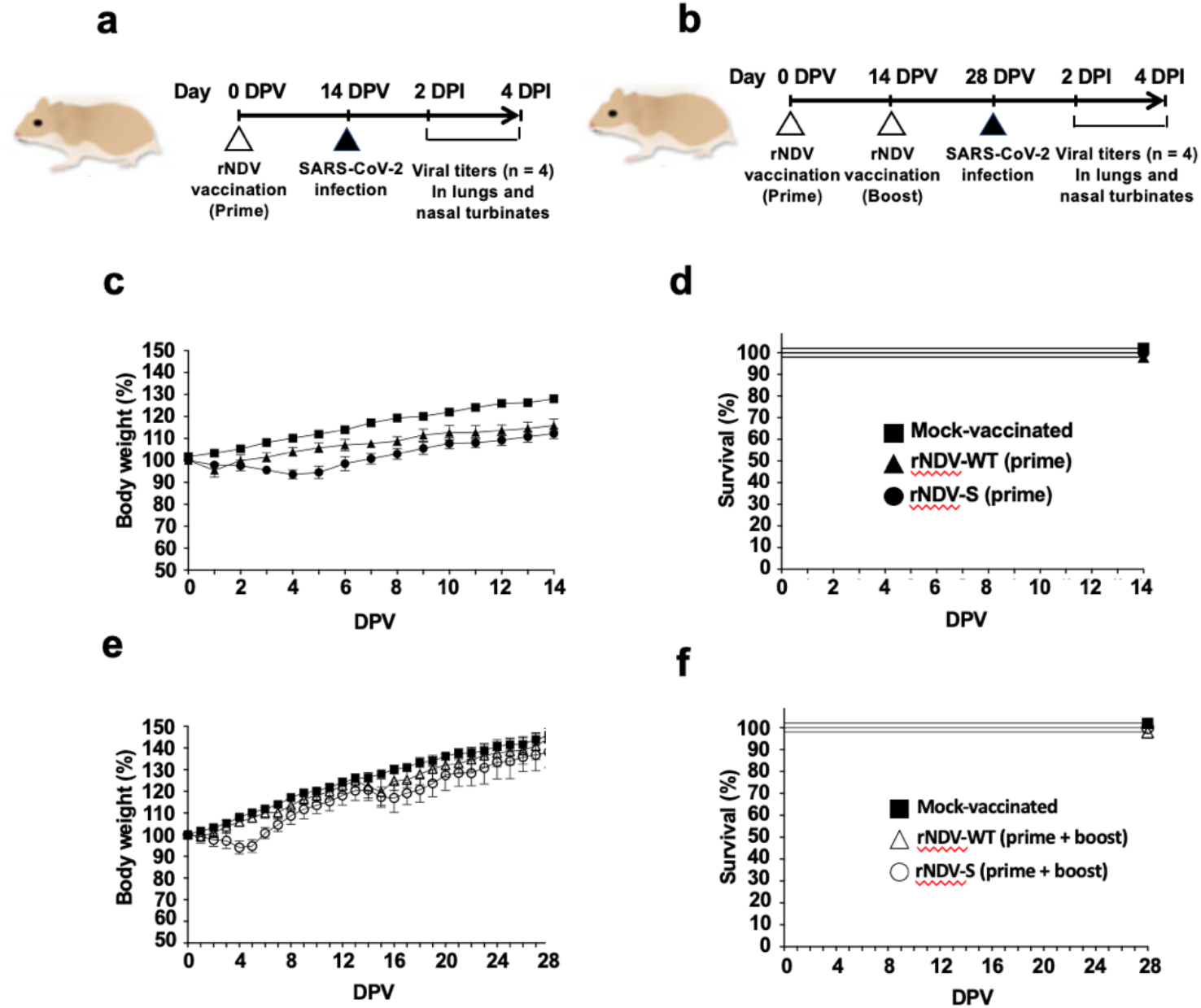
Safety of rNDV-S in golden Syrian hamsters. (a-b) Schematic representation of the vaccination approach: Golden Syrian hamsters (n=8/group) were mock (PBS)-vaccinated or vaccinated intranasally with 1 × 10^6^ PFU of rNDV-WT or rNDV-S using a prime (**a**) or a prime + boost (**b**) vaccination regimen. After vaccination, hamsters were challenged at 14 (prime, **a**) or 28 (prime + boost, **b**) DPV with 2 × 10^4^ PFU of SARS-CoV-2 and sacrificed at 2 and 4 DPV for gross lung pathology, viral titers, H&E staining and histopathology. (**c-d) Morbidity and mortality of rNDV-S in prime vaccinated hamsters:** Body weight changes (**c**) and survival (**d**) were evaluated at the indicated DPV with a single dose of rNDV-WT or rNDV-S. Mock (PBS)-vaccinated hamsters were used as control. (**e-f) Morbidity and mortality of rNDV-S in prime + boost vaccinated hamsters:** Body weight changes (**e**) and survival (**f**) were evaluated at the indicated DPV with two doses of rNDV-WT or rNDV-S. Mock (PBS)-vaccinated hamsters were included as control. Error bars represent standard deviations (SD) of the mean for each group of hamsters.

**Fig 6.**
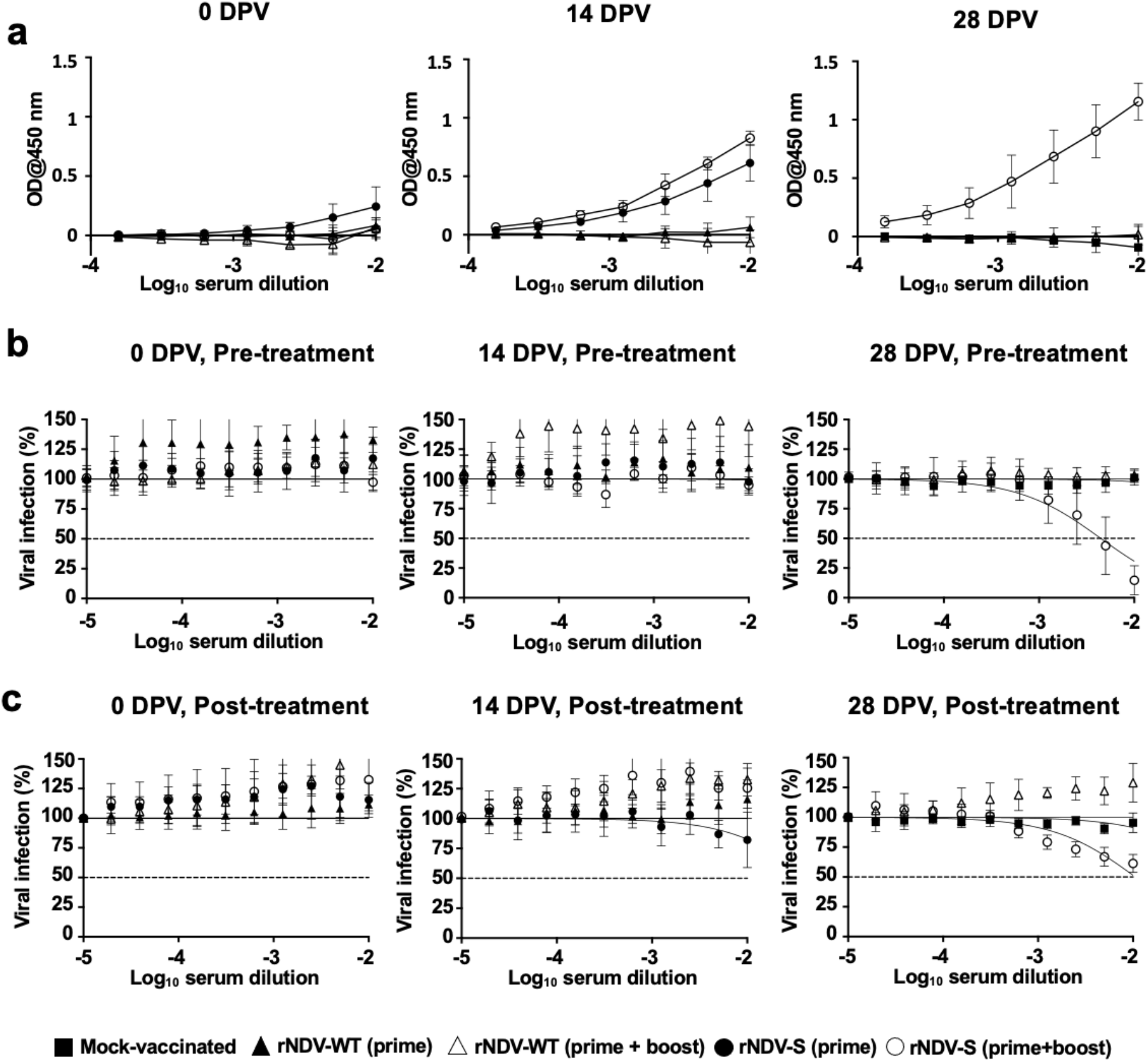
Total and NAbs in serum of golden Syrian hamsters vaccinated with rNDV-S. **(a) ELISA:** Levels of total S binding Abs in sera from hamsters vaccinated with rNDV-S at 0, 14, and 28 DPV. **(b) Neutralization assays, pre-treatment conditions:** *In vitro* neutralizing activity of hamster serum samples against SARS-CoV-2 at 0, 14, and 28 DPV. (**c) Neutralization assays, post-treatment conditions:** *In vitro* neutralizing activity of hamster sera against SARS-CoV-2 at 0, 14, and 28 DPV. Virus neutralization assays were quantified using ELISPOT and the percentage of infectivity calculated using sigmoidal dose response curves (**b and c**). Mock-infected cells and SARS-CoV-2 infections in the absence of serum were used as internal controls (**b and c**). Dotted line indicates 50% neutralization (**b and c**). Data were expressed as mean and SD.

### Safe intranasal administration of rNDV-S in hamsters

After establishing the basis of immunological and safety profiles in mice, we next attempted to evaluate the safety of aerosol rNDV-S vaccination in golden Syrian hamsters, which have the advantage of being susceptible to SARS-CoV-2 infection. A total of n=8 hamsters in each group were mock (PBS)-vaccinated or vaccinated with 1 × 10^6^ PFU of rNDV-WT or rNDV-S once (prime, Fig. 5a) or twice in a 2-week interval (boosted, Fig. 5b). Body weight was measured daily for 14 (prime) or 28 (boosted) DPV. Prime vaccination with rNDV-S resulted in no apparent clinical disease except slight (<5%) body weight loss at 5 DPV (Fig. 5c), and no mortality (Fig. 5d). Boosted vaccination also induced less than 5% of body weight loss by 5 DPV (Fig. 5e) with all the animals surviving rNDV-S vaccination (Fig. 5f). Vaccination with rNDV-WT resulted in body weight loss only one day after vaccination in either the prime or boosted vaccinated hamsters (Fig. 5c, e). These results demonstrate that vaccination of golden Syrian hamsters with 1 × 10^6^ PFU of rNDV-S is safe without lasting significant changes in body weight, with all the animals surviving aerosol rNDV-S vaccination.

### Ability of intranasally administered rNDV-S to induce Ab and NAb responses in hamsters

Serum samples collected at 0, 14, and 28 DPV from experimental hamsters were analysed for Ab and NAb responses against SARS-CoV-2. Total levels of Abs against SARS-CoV-2 were evaluated by ELISA, while NAbs were evaluated by PRNT assay using SARS-CoV-2. Sera from prime vaccinated hamsters (14 DPV) were able to react with extracts from SARS-CoV-2 infected cell homogenates, but not from mock-infected cell homogenates (Fig. 6a). Notably, levels of total Abs against SARS-CoV-2 were higher in sera from boosted rNDV-S-vaccinated hamsters (28 DPV) (Fig. 6a, Supplementary Fig. 7, 8). In terms of NAb responses, only sera from hamsters receiving boosted vaccination (28 DPV) presented NAbs in the PRNT assay in both pre-treatment (Fig. 6b, and Table 1) and post-treatment (Fig. 6c and Table 1) conditions. These results indicate that boosting is required to induce robust NAb responses against SARS-CoV-2 upon vaccination with rNDV-S.

**Fig 7.**
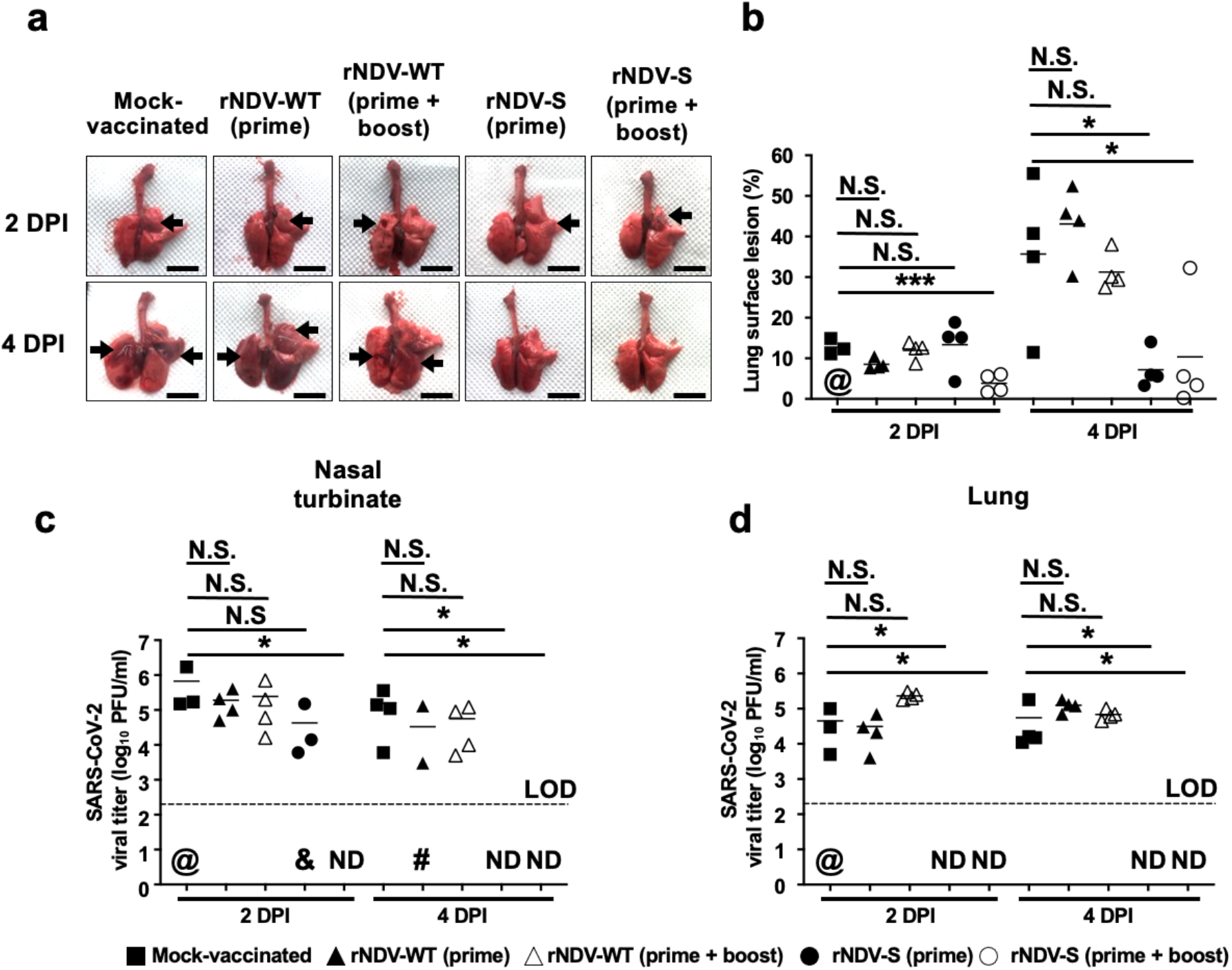
Gross lung pathology and SARS-CoV-2 titers in challenged golden Syrian hamsters. **(a) Lung images:** Representative images of the lungs from mock, rNDV-WT and rNDV-S vaccinated hamsters at 2 and 4 DPI with SARS-CoV-2 are shown. Scale bars = 1 cm. **(b) Lung pathologic lesions:** Macroscopic pathology scoring of lungs from hamsters in panel A was determined by measuring the distributions of pathological lesions (arrows), including consolidation, congestion, and pneumonic lesions using ImageJ software. (**c-d) SARS-CoV-2 titers:** SARS-CoV-2 titers in nasal turbinate (**c**) and lungs (**d**) from hamsters in panel A were determined by standard plaque assay. Dotted line indicates limit of detection (LOD, 200 PFU). Each symbol represents an individual animal and @ represents one mock-vaccinated hamster that was removed because of accidental death. &, Virus not detected in one mouse; #, virus not detected in two mice; ND, not detected. Lines represent the geometric mean. The Mann–Whitney test used for statistical analysis. *, P<0.05; **, P<0.01; or ***, P<0.005 for indicated comparisons.

**Fig 8.**
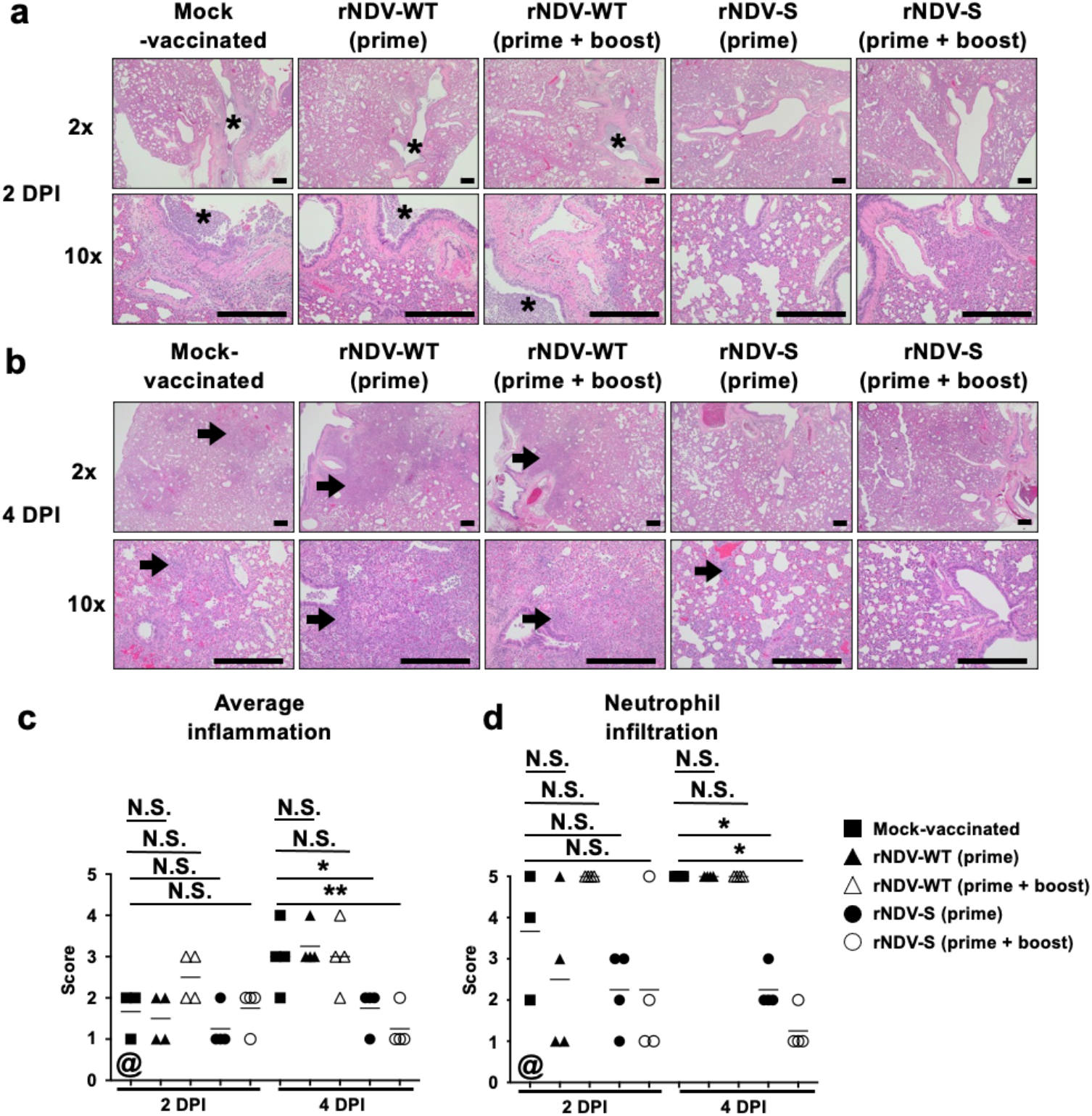
Histopathologic analysis of SARS-CoV-2 challenged golden Syrian hamsters. (a-b) H&E images of lungs from SARS-CoV-2 challenged hamsters at 2 DPI (a) and 4 DPI (b): Lung lesions are primarily characterized by suppurative inflammation within and surrounding the bronchi/bronchiole (black asterisks) at 2 DPI and extending to bronchointerstitial inflammation (arrows) by 4 DPI. Scale bars = 500 µm. **(c) Average inflammation scoring:** The average inflammation scores were determined based on percent of inflamed lung area: grade 0 = no histopathological lesions; grade 1= minimal (< 10%) histopathological lesions; grade 2 = mild (10 to 25%) histopathological lesions; grade 3 = moderate (25 to 50%) histopathological lesions; grade 4 = marked (50 to 75%) histopathological lesions; and grade 5 = severe (> 75%) histopathological lesions.**(d) Neutrophil infiltration:** The neutrophil infiltration scores were graded base on lesion severity as follows: grade 0 = no lesions observed; grade 1 = <10 cells; grade 2 = <25 cells; grade 3 = <50 cells; grade 4 = >100 cells; and grade 5 = too many cells to count. The Mann–Whitney test was used for statistical analysis. *, P<0.05; **, P<0.01; or ***, P<0.005 for indicated comparisons. Lines represent the geometric mean. @ represents one mock-vaccinated hamster that was removed because of accidental death.

### Protection efficacy of rNDV-S against SARS-CoV-2 infection in hamsters

To access protection efficacy of rNDV-S against SARS-CoV-2 infection, hamsters were vaccinated (prime or boosted) with rNDVs (WT or S) and then challenged with 2 × 10^4^ PFU of SARS-CoV-2. For viral titration and pathology, hamsters were sacrificed at 2 and 4 DPI (n=4/group). Mock-vaccinated hamsters either challenged with SARS-CoV-2 or mock challenged were used as internal controls. Lungs from mock-vaccinated hamsters showed mild to moderate, multifocal pneumonic lesions and congestions at 2 DPI, and higher inflammation scores characterized by moderate to severe locally extensive to diffuse bronchopneumonia and foamy exudate in the trachea at 4 DPI (Fig. 7a). Similar results were observed in hamsters vaccinated with rNDV-WT (primed or prime + boost). Conversely, hamsters vaccinated with rNDV-S (prime + boost) showed significantly lower inflammation scores compared to those of mock or rNDV-WT vaccinated hamsters at both 2 and 4 DPI (Fig. 7a, b).

Viral titers from nasal turbinate and lung followed similar trends and further support the lung inflammation scoring. We were not able to detect the presence of SARS-CoV-2 in the nasal turbinate and lungs of hamsters vaccinated with rNDV-S (boosted). Hamsters vaccinated with a single dose of rNDV-S (prime) showed SARS-CoV-2 titers in nasal turbinate and lungs at 2 DPI similar to those of mock-vaccinated hamsters or hamsters vaccinated with rNDV-WT (Fig. 7c, d). However, at 4 DPI, we could not detect the presence of SARS-CoV-2 in the nasal turbinate or lungs of hamsters vaccinated with rNDV-S, contrary to the situation of mock-vaccinated hamsters or hamsters vaccinated with rNDV-WT, where high titers of SARS-CoV-2 were detected at 4 DPI (Fig. 7c, d). Histopathological analysis further confirmed these results (Fig. 8). At 2 DPI, lungs from mock- or rNDV-WT-vaccinated hamsters showed abundant infiltration of inflammatory cells such as degenerate and non-degenerate neutrophils, macrophages, lymphocytes, plasma cells and few eosinophils within the lumen of bronchi and bronchioles and surrounding small blood vessels (Fig. 8a). At 4 DPI, histologic lesions were primarily characterized by extensive infiltration of the alveolar septa and alveolar space by neutrophils, macrophages and lesser lymphocytes, plasma cells, and few eosinophils. The bronchi and bronchioles showed a slightly lesser degree of inflammation compared to 2 DPI (Fig. 8b). No significant differences in inflammation (Fig. 8c) and neutrophil infiltration scores (Fig. 8d) were seen between the different groups at 2 DPI. However, degree of neutrophil infiltration and lung inflammation scores were significantly decreased in the rNDV-S vaccinated groups (inflammation score of 2.25 for prime and of 1.25 for boosted) compared to that of mock-vaccinated (inflammation score of 5) or infected with rNDV-WT (inflammation scores of 5 for both prime and boosted) groups at 4 DPI (Fig. 8d). Lungs of SARS-CoV-2 infected hamsters vaccinated with rNDV-S showed lower lung inflammation and numbers of neutrophil infiltration than those mock- or rNDV-WT vaccinated, indicating that a booster vaccination with rNDV-S protects hamsters from SARS-CoV-2 infection and subsequent lung damage (Fig. 8). Finally, IHC staining was performed to evaluate the presence and spread of SARS-CoV-2 replication by detecting viral N protein in the lungs from mock-, rNDV-WT-, or rNDV-S-vaccinated hamsters. As expected, SARS-CoV-2 N protein staining was not detected in any of the lungs from rNDV-S vaccinated groups (prime or boosted) at 2 and 4 DPI, while lungs from mock- or rNDV-WT-vaccinated hamsters showed significant SARS-CoV-2 N protein positively stained cells. At 2 DPI, SARS-CoV-2 viral N protein staining was primarily within the epithelium of bronchi, bronchioles, and in inflammatory cells present in the lumen of bronchi and bronchioles and extended mild to rarely to the alveolar septa surrounding the bronchioles. At 4 DPI, SARS-CoV-2 N protein positive straining was present in the epithelium of bronchioles, and along the edges of areas of bronchointerstitial inflammation within the inflammatory cells and alveolar septa that are distant from bronchioles (Fig. 9a). The average IHC scores showed progressive and widespread distribution of SARS-CoV-2 in mock- and rNDV-WT-vaccinated groups and marked absence of viral N protein staining at 2 and 4 DPI in the rNDV-S-vaccinated groups (Fig. 9b). These results demonstrate that vaccination with rNDV-S can protect from SARS-CoV-2 infection, resulting in lower SARS-CoV-2 titers and protection from lung damage.

**Fig 9.**
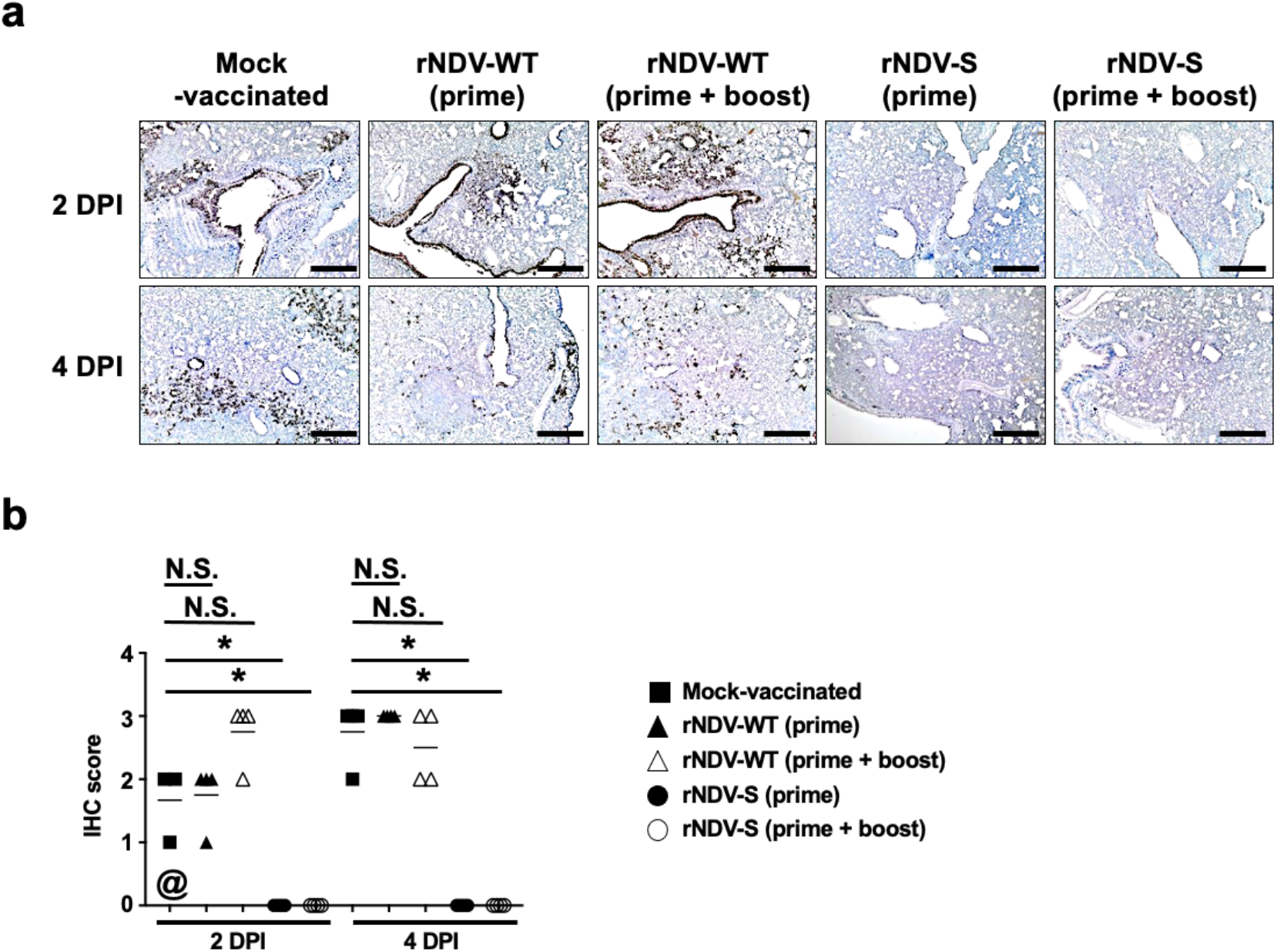
IHC analysis in SARS-CoV-2 challenged golden Syrian hamsters. **(a) IHC imaging:** Presence of SARS-CoV-2 viral N protein in the lungs of SARS-CoV-2 challenged hamsters at 2 and 4 DPI. Scale bars = 500 µm. **(b) IHC scores:** Scores were determined on the presence of SARS-CoV-2 N protein staining: grade 0 = no or rare immunostaining; grade 1 = viral N protein staining only in bronchi/bronchiole; grade 2 = viral N protein staining in bronchi/bronchiole and surrounding alveolar septa; grade 3 = viral N protein staining in alveolar septa distant from the bronchi and bronchioles; grade 4 = viral N protein staining throughout the lung. @ represents one mock-vaccinated hamster that was removed because of accidental death. The Mann–Whitney test was used for statistical analysis. *, P<0.05; **, P<0.01; or ***, P<0.005 for indicated comparisons.

## DISCUSSION

Due to high transmissibility of SARS-CoV-2 and lack of considerable pre-existing immunity, the COVID-19 pandemic is posing significant health challenges particularly in the elderly and people with existing co-morbidities^37-39^. While both treatments and vaccines are being developed^40-44^, there are limited attempts to offer vaccines that can be deployed in frail and resource-limited health care system particularly in low- and middle-income countries. In a thirst to curb the pandemic, upper and middle-income countries are scooping most of the pharmaceutical capacity of vaccine production, which can contribute to reduce the transmission and impact of the pandemic. In this regard, a low-cost vaccine production system for mass immunization of people in low and middle-income countries is required. Here, we describe a promising and scalable live-attenuated vector vaccine based on the use of a rNDV expressing SARS-CoV-2 S, which can be produced robustly and economically in avian embryonated eggs, as well as in US FDA-approved cell lines, that can be administered intranasally to induce protection at the site of SARS-CoV-2 infection.

Our studies establish that intranasal vaccination with rNDV-S induces Abs, including NAbs, against SARS-CoV-2 full S protein and RBD, as well as antigen specific T-cell responses. Although a single dose confers protection against SARS-CoV-2 challenge, a prime and boost regime provided superior protection, reduced pathology in lung and completely blocked shedding of SARS-CoV-2 in hamsters, which is considered an appropriate animal model for vaccine evaluation^45,46^. The intranasal route for immunization is advantageous in comparison to traditional routes (i.e., intramuscular) mainly due to its ability to elicit immunity at the local mucosal level^47-49^. Enhancing mucosal protective abilities is desirable as it is the first line of defence in the human body against the majority of pathogens targeting the respiratory tract. Beside the possibility to self-administer and lack of needles, the intranasal delivery of antigen is associated with mucosal immunity including production of IgA and priming of T and B cells in the nasopharynx-associated lymphoid tissues^50^, and may also provide cross-protection by eliciting mucosal immunity at different mucosal sites such as the intestines and genital tract, which have been shown to be potential replication sites for SARS-CoV-2^51,52^. Indeed, intranasal vaccination of mice with rNDV-S induced the secretion of IgA as well as IgG2a Abs providing a complete response both locally and systemically. Moreover, detection of significant increase in Abs in pleural lavage fluid highlight the local induction of the immune response post-intranasal vaccination. While significant Abs were elicited, further increasing the interval between prime and boost doses may enhance the memory responses and higher humoral immunity which warrant future studies. Nevertheless, the induction of Abs following administration of intranasal vaccines does not require highly skilled personnel and could even be self-administered, while also avoiding the issues of individuals who experience trypanophobia and hemophobia. Based on these advantages, we propose that intranasal delivery of rNDV-S is a promising platform for preventing SARS-CoV-2 infection, COVID-19 disease, and transmission and, therefore; it warrants clinical evaluation in humans.

The correlation between SARS-CoV-2 viral load, viral manifestation, clinical symptoms and the variety of immune cells associated with viral immunity are a topic of ongoing debate. However, the relation between the site of cellular infiltration and kinetics of sensitive cell/humoral immunity is by far the most influential factor determining the outcome of the infection. CD8+ T cells are critical in the fight against viral infections as their cytotoxic characteristics are designed to target intracellular threats such as viruses. CD4+ T cells on the other hand drive the immune milieu towards a Th1 immune response characterised by IFNγ-mediated immune responses and subsequent IgG2a Ab release. Current evidence indicate that Th1 type responses are key in the successful control of SARS-CoV-2^53^. In our study, systemically, spleen CD4+ T cell IFNγ^+^ levels increased post-vaccination in both prime and boost groups. Locally, in the lung tissue, increased IFNγ levels from both CD4+/CD8+ T cells were observed. This, in line with previously discussed data potentially provides a level of protective immunity in the form of IFNγ-antiviral functions^45^. Boosts in TNF have been associated with disease severity. TNF and IL-10 directly correlate with COVID-19 severity and are synonymous with patient deterioration (SARS-CoV-2: A Storm is Raging)^54^. In SARS-CoV-2 infection, TNF appears to lean towards detrimental rather than protective roles, especially as a key component of the group of cytokines known as the cytokine storm^55^. In our work, a reduction in lung CD8+ T cell-TNF^+^ levels were associated with increased CD8+ T cell percentages. This potentially suggests that in our model, the vaccine promotes cytotoxic phenotype for CD8+ T cells independent of TNF.

Overall induction of IgM, IgA and IgG as well as NAbs have been reported in numerous natural infections and in vaccine trials^13^. Coronavirus-specific IgM production is transient and leads to isotype switch to IgA and IgG, these latter Ab subtypes can persist for extended periods in the serum and in nasal fluids^45^. After two doses of intranasal vaccination, we hypothesized the greater protection observed was because of the mucosal immune responses generated. In our study, at day 19 post immunisation, local IgM levels were not increased in either FALC or plural fluid suggesting a natural transition of the Ab phenotype towards an adaptive response from an innate immediate reaction. Interestingly, studies have reported earlier peaks in anti-S protein IgA in mild cases earlier than peaks in IgM^54,56^. This suggests a more robust effect for IgA in COVID-19 in protection and clinical outcome of the disease^57^. Indeed, high levels of anti-SARS-CoV-2 IgA were detected in serum and lung, and B cells secreting IgA were detected in the spleen only in mice vaccinated via an intranasal route. Moreover, intranasal but not intramuscular vaccination induced SARS-CoV-2-specific CD8+ T cells in the lung, including CD103+CD69+ cells, which are likely of a resident memory phenotype^13^. Future Ab passive transfer and T cell depletion studies can assess the relative contribution of each arm of the immune system and establish more precisely the mechanistic basis for the enhanced protection conferred by intranasal delivery of rNDV-S vaccine.

In our study, we noted an increase in specific anti-S RBD-IgA levels in the pleural cavity. This corroborated with previous findings associating with mild COVID-19 infections where anti-S RBD-IgA levels were elevated at different mucosal tissues in recovered COVID-19 patients^57,58^. Conversely, in our study, anti-S RBD-IgG2a production increased in the pleural cavity. This localised expansion of IgG2a, in particular, suggests a promotion of an Fc-mediated killing of the target cell through antibody-dependent cellular cytotoxicity (ADCC) type of protection in response to the vaccine. In order to assess the relative contribution of each arm of the immune system to conferred protection against SARS-CoV-2 by intranasal delivery of rNDV-S vaccine, future studies are warranted.

To our knowledge, there are limited, if any, SARS-CoV-2 vaccine platforms currently in clinical trial using intranasal delivery methods. Similar to influenza LAV, which induces profound local humoral and cellular responses^59^, immunization with rNDV-S offer superior properties over the intramuscular route. Additionally, sterilizing immunity against re-infection or blockage of virus shedding require local and mucosal immunity against SARS-CoV-2, which is optimally provided by the intranasal route^60,61^.

In summary, our studies established that intranasal immunization with rNDV-S induces both NAbs and antigen-specific T cell responses in mice. Intranasal administration of two doses fully protected hamsters against lung infection, inflammation, and pathology after SARS-CoV-2 challenge. Importantly, single or double dose of rNDV-S completely blocked SARS-CoV-2 shedding in the nasal turbinate and lungs of hamsters. Thus, intranasal vaccination with rNDV-S has the potential to control SARS-CoV-2 infection at the site of inoculation, which should prevent both virus-induced COVID-19 disease and transmission. Based on these pre-clinical results in two rodent models, future clinical evaluation of rNDV-S in humans for the treatment of SARS-CoV-2 infection is urgently warranted.

## METHODS

### Animals

BALB/c female mice were purchased from Charles River, UK and housed under specific pathogen free conditions at Lancaster University. Experiments were performed using age and sex matched mice at 11-14 weeks of age. Mice were provided sterilized water and chow *ad libitum* and acclimatized for minimum of one week prior to experimental manipulation. Specific-pathogen-free (SPF) 4-weeks-old female golden Syrian hamsters were purchased from Charles River, US. Hamsters were maintained in micro-isolator cages at Animal Biosafety Level (ABSL)-2 before and during vaccination with rNDV-WT or rNDV-S. For challenge with SARS-CoV-2, hamsters were transferred to ABSL-3. Hamsters were provided sterilize water and chow *ad libitum* and acclimatized for one week prior to experimental manipulation. All experiments in hamsters were conducted at Texas Biomedical Research Institute.

### Ethics statement

All mice experiments were performed under the regulations of the Home Office Scientific Procedures Act (1986), specifically under the project licence PPL PACEB18AC. The project licence was approved by both the Home Office and the local AWERB committee of Lancaster University. All the *in vitro* and *in vivo* hamster experiments with infectious SARS-CoV-2 were conducted under appropriate BSL-3 and ABSL-3 laboratories, respectively, at Texas Biomedical Research Institute, San Antonio, Texas, US. Experiments were approved by the Texas Biomedical Research Institutional Biosafety (BSC) and Institutional Animal Care and Use (IACUC) committees, protocols BSC20- and 1722 MA 3, respectively.

### Cells and viruses

SARS-CoV-2, USA-WA1/2020 strain (Gen Bank: MN985325), was obtained from the Biodefense and Emerging Infections Research Resources Repository (BEI Resources, NR-52281) and amplified in African green monkey kidney epithelial Vero E6 cells obtained from the American Type Culture Collection (ATCC, CRL-1586) as previously described^62^. Briefly, the virus stock obtained from BEI Resources considered as passage four (P4) was amplified two more times to generate a P6 working stocks for animal infections. To that end, Vero E6 cells were infected at low multiplicity of infection (MOI, 0.01) for 72 h and tissue culture supernatants (TCS) were collected, clarified, aliquoted and stored at −80°C until use. Virus stocks were titrated in Vero E6 cells by plaque assay and immunostaining as previously described^35,63^. The rNDV-WT and rNDV-S viruses were propagated in chicken embryonated eggs quantified in Vero cells using procedures we described before^33^. 293T cells and Vero cells were maintained in Dulbecco’s Modified Eagle Medium (DMEM) with 10% FBS as described previously^33^.

### Safety and immunogenicity studies in mice

Mice were assigned to 4 experimental groups receiving a 50 µl intranasal dose of either mock (PBS) vehicle at days 0 and 14; 10^4^ PFU of rNDV-WT at days 0 and 14 (booster) or rNDV-S at day 0 only (prime) or at days 0 and 14 (booster). Mice and chow were weighed daily and nasally swabbed at times indicated before sacrifice at 19 DPV. Mouse morbidity was scored using a 14 point scoring system based on a) Hunched posture: Yes (1), No (0); b) Spontaneous activity: None (2), Reduced (1), Normal (0); c) Response to touch: None (3), Movement (2), Move away (1), Normal (0); d) Feels cold: Yes (1), No (0); e) Breathing laboured: Yes (1), No (0); f) “Rasping” breathing: Yes (1), No (0); g) Ruffled fur/piloerection: Yes (1), No (0); h) Pallor at extremities: Yes (1), No (0); and i) Moderate staining around eyes and/or nose.

### Vaccine safety studies in hamsters

To evaluate *in vivo* safety of rNDV-S, hamsters (n=8 per group) were vaccinated intranasally with 1 × 10^6^ PFU of rNDV-WT or rNDV-S, or mock-vaccinated with saline (PBS), in a final volume of 100 µl, after sedation in an isoflurane chamber. Hamsters were vaccinated following a prime or a booster regimen. In the case of the booster immunization, animals were boosted 2 weeks after prime vaccination. After vaccination, hamsters were monitored daily for morbidity (body weight changes and clinical signs of infection) and mortality (survival) for 14 (prime) or 28 (booster) DPV. Sera were collected at 0 and 14 (mock, prime, and booster groups), and 28 (booster) DPV. One mock-vaccinated hamster in the vaccination group was removed because of accidental death.

### Vaccine efficacy in hamster

After 14 (prime) or 28 (mock and booster groups) DPV, hamsters were challenged intranasally with 2 × 10^4^ PFU of SARS-CoV-2 in a final volume of 100 µl under isoflurane sedation. To evaluate SARS-CoV-2 titers in nasal turbinates and lung, hamsters were humanely sacrificed at 2 (n=4) or 4 (n=4) DPI. Nasal turbinates and lungs were harvested, and half of the organs were homogenized in 2 mL of PBS using a Precellys tissue homogenizer (Bertin Instruments) for viral titration and the other half was kept in 10% neutral buffered formalin (NBF, ThermoFisher Scientific) for histopathology and immunohistochemistry (IHC). Tissue homogenates were centrifuged at 21,500 x *g* for 5 min and supernatants were used to calculate viral titers.

### Real-time PCR for quantification of rNDV replication

RNA was extracted from mice organs (nasal turbinate, lungs, trachea, and gut) using TRIzol™ reagent as per manufacturer’s instructions (Invitrogen, USA). Real-time qRT-PCR was performed using SuperScript™ III Platinum™ One-Step qRT-PCR Kit (Invitrogen, USA) following the manufacturer’s instructions to detect NDV M gene^64^ and enable the calculation of NDV genome copies.

### Production and purification of recombinant SARS-CoV-2 S and RBD proteins

Recombinant S and RBD proteins were produced and purified as previously described^65^. Briefly, expression cassettes containing SARS-CoV-2 S and RBD nucleotide sequences were codon optimized for Drosophila melanogaster Schneider 2 (S2) cells. The N-terminus signal sequence of both S and RBD proteins were replaced with Drosophila BiP signal sequence which encodes immunoglobulin-binding chaperone protein, and the C-terminus were fused with T4 foldon sequence (GSG YIP EAP RDG QAY VRK DGE WVL LST FL) and a C-tag sequence (EPEA) for affinity purification. The protein expression cassettes were commercially synthesised (GeneArt, ThermoFisher Scientific, Regensburg, Germany) and cloned into pExpres2.1 expression vector (ExpreS2ion Biotechnologies, Hørsholm, Denmark). The recombinant plasmids were transfected into Drosophila S2 cells using Calcium Phosphate Transfection Kit (Thermo Fisher Scientific, Paisley, Scotland, UK). Following antibiotic selection with Zeocin (InvivoGen, Toulouse, France), cells were propagated in Drosophila EX-CELL® 420 Serum-Free Medium (Merck Life Science) at 25°C. Recombinant S and RBD trimeric proteins secreted in cell culture supernatants were purified using the CaptureSelect™ C-tag Affinity Matrix (ThermoFisher Scientific). Concentration of purified recombinant Abs were determined by PierceTM BCA Protein Assay Kit (ThermoFisher Scientific) and the purity was assessed by sodium dodecyl sulphate–polyacrylamide gel electrophoresis (SDS-PAGE).

### Indirect antigen capture enzyme-linked immunosorbent assays (ELISA) for Ab detection in mice

SARS-CoV-2 S or RBD proteins were coated onto 96-well ELISA plates. Viral antigen proteins were diluted in PBS to a concentration of 5 µg/ml, using 100 µl per well. Following an overnight incubation at 4°C, plates were washed 3 times with PBS-Tween-20 (PBST) and blocked with 2% bovine serum albumin (BSA) in PBS overnight at 4°C. Sera samples were heat inactivated at 56°C for 30 min prior to testing by ELISA. Hundred µl of the diluted (1:10 in 2% BSA) primary Abs were added to each well and incubated for 3 h at 37°C. Following 3 times washing with PBST, plates were incubated with 1:6,000 IgG1 HRP avidin-horseradish-peroxidase conjugate diluted in 2% BSA for 2 h at 37°C. Following 3 times washing in PBST, colour was developed using 3,3’,5,5’-tetramethylbenzidine (TMB) chromogen (100 µl per well) for 5 min at room temperature. The reaction was stopped with an equal volume of 0.16M H_2_SO_4_ and optical density (OD) was read at 450 nm. Sample to positive ratio (S/P), as defined by the formula S/P = (sample OD - standard negative OD)/ (standard positive OD - standard negative OD), were used for reporting all ELISA values.

### ELISA for Ab detection in hamsters

Binding affinity of hamster sera to SARS-CoV-2 antigens was determined by ELISA using lysates from mock-vaccinated and SARS-CoV-2-infected Vero E6 cells. Briefly, ELISA plates were coated overnight at 4°C with a 1:1,000 dilution of cells extracts from mock- and SARS-CoV-2-infected Vero E6 cells. Plates were washed 3X with PBS and incubated with 2-fold, serially diluted hamster serum samples in PBS (starting dilution of 1:100) for 1 h at 37°C. Plates were washed 3X with PBS and incubated with HRP-conjugated anti-hamster IgG (Jackson ImmunoResearch, PA, USA) for 1 h at 37°C. After 3X washes with PBS, plates were developed with TMB substrate buffer. The reaction was stopped by 2M H_2_SO_4_ after incubation for 10 min at room temperature. The optical density (OD) values were measured at 450 nm using an ELISA plate reader.

### Neutralization assays using pseudotyped viruses

Derivatives of 293T cells expressing ACE2 receptors were generated by transducing 293T cells with ACE2 receptor expressing vector. Human 293T cells expressing ACE2 were used for rescue of SARS-COV-2 S-pseudotyped HIV particles for neutralisation assays. To generate SARS-COV-2 S-pseudotyped HIV, the reporter vector (pCCNanoLuc2AEGFP), HIV-1 structural/regulatory proteins (pHIVNLGagPol) and pSARS-CoV-2-Strunc were used at a molar plasmid ratio of 1:1:0.45 and the infectivity of pseudotyped viral particles was calculated as described previously^34^. For neutralization assays of mice sera, sera (1:10 starting dilution) were five-fold serially diluted in 96-well plates. Thereafter, a 50 μl aliquot of HIV-based SARS-CoV-2 pseudovirus containing approximately 1×10^3^ infectious units were added. After 1h incubation at 37°C, 100 μl of the mixture was transferred to target cells plated at 1×10^4^ cells/well in 100 μl medium in 96-well plates. Cells were then cultured for 48h and cells were harvested for NanoLuc luciferase assays. For the NanoLuc luciferase assays, cells were washed twice, carefully, with PBS and lysed with 50 μl/well of Luciferase Cell Culture Lysis reagent (Promega). NanoLuc Luciferase activity in lysates was measured using the Nano-Glo Luciferase Assay System (Promega). Specifically, 25 μl of substrate in NanoGlo buffer was mixed with 25 μl cell lysate in black flat bottom plates and incubated for 5 min at RT. NanoLuc luciferase activity was measured using Glowmax Navigator luminometer (Promega), using 0.1s integration time. Relative luminescence units (RLU) obtained were normalized to those derived from cells infected with SARS-CoV-2 pseudovirus in the absence of Abs. The half maximal neutralizing concentration (NC_50_) for sera was determined using 4-parameter nonlinear regression curve fit to raw infectivity data measured as relative light units, or as the percentage of infected cells (GraphPad Prism).

### SARS-CoV-2 microneutralization assay

The day before infection, Vero-E6 cells were seeded at 1×10^4^ cells/well into 96-well plates and incubated at 37°C in a 5% CO_2_ incubator overnight. Mice sera samples were heat inactivated at 56°C for 30 min. Sera samples were 10-fold serially diluted in DMEM (Invitrogen, USA) supplemented with 1% BSA and 1x penicillin/streptomycin. Next, the diluted sera samples were mixed with a constant amount of SARS-CoV-2 (100 Median Tissue Culture Infectious Dose (TCID_50_)/50 μl) and incubated for 1 h at 37°C. The Ab-virus-mix was then directly transferred to Vero E6 cells and incubated for 3 days at 37°C in a 5% CO_2_ incubator. The median microneutralization (MN) 50 (MN_50_) or 100 (MN_100_) of each serum sample was calculated as the highest serum dilution that completely protect the cells from cytopathic effect (CPE) in half or all wells, respectively.

### Plaque reduction microneutralization (PRMN) assays

PRMN assays were performed to identify the levels of SARS-CoV-2 NAbs as described previously^30^. Briefly, sera samples were inactivated at 56°C for 30 min prior to assay. For pre-treatment conditions, 200 PFU/well of SARS-CoV-2 were mixed with 2-fold serial dilutions (starting dilution 1:100) of hamsters’ sera and incubated at 37°C for 1 h. Confluent monolayers of Vero E6 cells (96-well plate format, 4 × 10^4^ cells/well, quadruplicates) were infected with the virus-serum mixture for 1 h 37°C. After viral absorption, infectious media was exchanged with post-infection media containing 2% FBS and 1% Avicel. For post-treatment conditions, confluent monolayers of Vero E6 cells (96-well plate format, 4 × 10^4^ cells/well, quadruplicates) were infected with 100 PFU/well of SARS-CoV-2 for 1 h at 37°C. After viral adsorption, infection media was exchanged with 100 µl of post-infection media containing 2% FBS, 1% Avicel, and 2-fold serial dilutions (starting dilution 1:100) of hamster sera. In both cases (pre- and post-treatment conditions) infected cells were fixed 24 h post-infection (hpi) with 10% neutral formalin for 24 h and immunostained with a monoclonal Ab (1C7) against the viral nucleocapsid (N) protein (1 µg/ml). Viral neutralization was evaluated and quantified using ELISPOT, and a sigmoidal dose-response, non-linear regression curve was generated using GraphPad Prism to calculate median neutralization titer (NT_50_) in each of the serum samples.

### SARS-CoV-2 virus titrations

Nasal turbinate and lungs from SARS-CoV-2-infected golden Syrian hamsters were homogenized in 2 ml of PBS for 20 s at 7,000 rpm using a Precellys tissue homogenizer (Bertin Instruments). Confluent monolayers of Vero E6 cells (96-plate format, 4 × 10^4^cells/well, duplicates) were infected with 10-fold serial dilutions of supernatants obtained from the nasal turbinate or lung homogenates from SARS-CoV-2 infected golden Syrian hamsters. Virus from serially diluted samples was adsorbed at 37°C for 1 h followed by incubation in post-infection media containing 2% FBS and 1% Avicel at 37°C for 24 h. After viral infection, plates were submerged in 10% NBF for 24 h for fixation/inactivation. For immunostaining, cells were washed 3X with PBS and permeabilised with 0.5% Triton X-100 for 10 min at room temperature, followed by blocking with 2.5% bovine serum albumin (BSA) in PBS for 1 h at 37°C. Cells were incubated with the N protein 1C7 monoclonal Ab (1 µg/ml) diluted in 1% BSA for 1 h at 37°C. Then cells were washed 3X with PBS and stained with the Vectastain ABC kit and developed using the DAB Peroxidase Substrate kit (Vector Laboratory, Inc, CA, USA), as previously described ^1^. Virus titers were calculated as PFU/ml.

### Cell re-stimulation and flow cytometry staining

Spleens were removed from mice and disaggregated through a 100 µm sieve. Pericardium was digested using 1mg/ml Collagenase D (Roche) in a shaking heat block at 37°C, digestion was stopped after 35mins by addition of 5mM EDTA (Fisher) followed by passing through a 100µm cell strainer. Pleural exudate cells (PLEC) were isolated via lavage of the pleural cavity with 10ml of ice cold dPBS (Sigma). Left lung lobes were excised and finely diced using scissors prior to digestion in 2 ml lung digest medium (PBS (Sigma), 0.1mg/ml Liberase TM (Roche), 50 µg/ml DNAse I (Roche)) for 30 min in a shaking incubator at 37°C. Digestion was stopped after 30mins by adding 5mM EDTA (Fisher) followed by passing through a 100 µm cell strainer. Lung & spleen cells were diluted to 1×10^6^cells/ml before incubating with 50 μg/ml SARS-CoV-2 purified spike antigen for 24 h in media (RPMI-1640, 10% FCS, 100U/ml penicillin/streptomycin, 5%NEAA, L-glutamine and HEPES, 0.05 mM β-mercaptoethanol (SIGMA)). Cell suspensions were incubated for the final 6 h with 1x Cell stimulation cocktail (plus protein inhibitors) (eBioscience). Cell suspensions were blocked with anti-FcγR Ab (clone 24G2; eBioscience and Biolegend) before labelling with combinations of Abs specific for Ly6G (1A8; Biolegend), Siglec F (E50-2440; BD), CD19 (6D5, Biolegend), TCRβ (H57-597, Biolegend), CD11c (N418; Biolegend), CD45 (30-F11; Biolegend), CD5 (53-7.3; Biolegend), Ly6C (HK1.4; Invitrogen), CD11b (M1/70; Biolegend), IA/IE (M5/114.15.2, Biolegend), F4/80 (BM8, Invitrogen), CD3 (eBio500A2), CD4 (clone GK1.5; eBioscience, RM4-5, Biolegend),, CD49b (DX5; EBioscience), IL-13 (clone eBiol13A; eBioscience), IFNγ (clone XMG1.2; eBioscience), Ki67 (REA183, Miltenyi) and IL-17(eBio17B7; eBioscience). For intracellular cytokine analysis cells were then stained with Abs using the eBioscience Foxp3 permeabilization kit according to the manufacturer’s instructions. All samples were analysed on a Beckman Cytoflex and analysed with FlowJo Software (TreeStar).

### Pathologic observation

Mouse lung tissues were inflated and fixed in 10% neutral buffered formalin (NBF) solution and embedded in paraffin prior to H&E. After mounting, positive cells were enumerated in field of view and alveolar space enumerated via NIH Image J software. In the hamster studies, lungs were collected at 2- and 4-DPI from mock- and SARS-CoV-2-challenged golden Syrian hamsters. Lung samples were photographed to show gross lesions on both the dorsal and ventral views. Images were used for macroscopic pathology scoring analysis by measuring the distributions of pathological lesions, including consolidation, congestion, and pneumonic lesions using NIH ImageJ software. The area of pathological lesions was converted into percent of the total lung surface area. Half of the hamster lungs fixed with 10% neutral buffer formalin (NBF) were embedded in paraffin blocks, and sectioned (5 µm). Sections were stained with H&E and evaluated under light microscopy in a blinded manner by a board-certified veterinary pathologist at Texas Biomedical Research Institute. The average inflammation scores were graded based on percent of inflamed lung area: grade 0 = no histopathological lesions observed, grade 1 = minimal (< 10%), grade 2 = mild (10 to 25%), grade 3 = moderate (25 to 50%), grade 4 = marked (50 to 75%), and grade 5 = severe (> 75%). And the neutrophil infiltration scores were graded base on lesion severity as follows: grade 0 = no lesions observed, grade 1 = <10 cells, grade 2 = <25 cells, grade 3 = <50 cells, grade 4 = >100 cells, and grade 5 = Too numerous to count (TNTC).

### Immunohistochemistry (IHC) assays

Immunostainings were performed as previously described^54,55^. Briefly, 5 um lung tissues sections were mounted on Superfrost Plus Microscope slides, deparaffinized and antigen retrieval was conducted using the HIER method (Heat Induced Epitope Retrieval).

Subsequently, slides were stained with the primary N protein monoclonal Ab 1C7 (1 µg/ml). Then slides were incubated and developed with the MACH 4 Universal HRP Polymer Kit and Betazoid DAB Chromagen Kit (Biocare Medical, LLC), according to the manufacturer’s instructions. The presence of SARS-CoV-2 N protein antigen was analysed by a veterinary pathologist at Texas Biomedical Research Institute. Scores of antigens were graded based on the anatomical location of viral N protein staining: grade 0 = no or rare immunostaining, grade 1 = viral N protein staining only in bronchi/bronchiole, grade 2 = viral N protein staining only in bronchi/bronchiole and surrounding alveolar septa, grade 3 = viral N protein staining in alveolar septa distant from the bronchi and bronchioles, grade 4 = viral N protein staining throughout the lung. The Mann–Whitney test was used for statistical analysis.

### Confocal imaging

Mouse mediastinum samples were fixed for one hour on ice in 10% NBF (Sigma) and then permeabilised in PBS 1% Triton-X 100 (Sigma) for 15 min at room temperature prior to staining with primary Abs for one hour at room temperature in PBS 0.5% BSA 0.5% Triton. DAPI was added for the final 10 min of the 1 h incubation. After washing in PBS, tissues were mounted in fluoromount G and confocal images were acquired using a Zeiss LSM880 confocal laser scanning microscope. Samples were analysed using Fiji software. Cluster area was delineated using IgM, and percentage area of Ki67 & CD4 staining within each cluster was calculated.

### Statistical analysis

Results are expressed as mean ± SEM. Where statistics are quoted, two experimental groups were compared via the Student’s t test or the Mann–Whitney for non-parametric data. Three or more groups were compared with ANOVA, with Sidaks, Dunnett’s or Bonferroni’s post-test as indicated. A P value of <0.05 was considered statistically significant. *, P<0.05; **, P<0.01; or ***, P<0.005 for indicated comparisons, error bars represent standard error of means.

## ACKNOWLEDGEMENTS

We want to thank Dr. Thomas Moran at the Icahn School of Medicine at Mount Sinai for providing us with the SARS-CoV cross-reactive N protein monoclonal Ab 1C7. We also thank BEI Resources for providing the SARS-CoV-2 USA-WA1/2020 isolate (NR-52281). We would also like to thank members at our institutes for their efforts in keeping them fully operational during the COVID-19 pandemic, and the BSC and IACUC committees at Texas Biomedical Research Institute for reviewing our protocols in a time efficient manner. We would also like to thank the support of Texas Biomed Research Institute members and donors, whose efforts and COVID-19 philanthropic donations, respectively, made possible the realization of this study. The MM laboratory is supported by the Biotechnology and Biological Sciences Research Council (BBSRC) (BB/M008681/1 and BBS/E/I/00001852) and the British Council (172710323 and 332228521). JJW was supported by a Wellcome Trust Grant; Award Number: 209087/Z/17/Z. LHJJ was supported by a Wellcome Trust Grant; Award Number 213697/Z/18/Z.

## AUTHOR CONTRIBUTIONS

The overall design of the study was by M.M., L.M., L.H.J. and J.J.W. Experiments and analyses were performed by J.P., F.S.O., M.A.R., M.M., J.W., J.T., L.H.J. and J.J.W. Recombinant protein for mice experiment was produced by M.I. and P.C. Animal studies were performed by L.H.J., J.J.W., L.M., J.B.T., V.S., R.E., J.P., F.S.O., J.W. and J.T. The manuscript was written by M.M., L.H.J. and J.J.W, J.P., and F.S.O. and revised by M.A.R, B.M.A., W.A., A.A., M.I., R.E., V.S., J.B.T., and L.M. All authors have read and approved the manuscript.

## ADDITIONAL INFORMATION

Supplementary information is available for this paper.

Correspondence and requests for materials should be addressed to MM.

