## Supplementary Information for "Immunogenicity and Protective Efficacy of an Intranasal Live-attenuated Vaccine Against SARS-CoV-2 in Preclinical Animal Models"

Article

*[1]Texas Biomedical Research Institute, Host-Pathogen Interactions and Population Health Programs, San Antonio, TX, 78227, USA; [2]Division of Biomedical and Life Sciences, Faculty of Health and Medicine, Lancaster University, Lancaster LA1 4YG, UK; [3]Department of Virology, Faculty of Veterinary Medicine, Cairo University, Giza, 12211, Egypt; [4]Faculty of Applied Medical Sciences, Department of Medical Laboratory Technology, Immunology Group, King Abdul Aziz University, Jeddah, Saudi Arabia; [5]Faculty of Applied Medical Sciences, Department of Medical Laboratory Technology, Microbiology Group, King Abdul Aziz University, Jeddah, Saudi Arabia; [6]Faculty of Medicine, Al Baha University, Al Baha, Saudi Arabia; [7]The Pirbright Institute, United Kingdom*

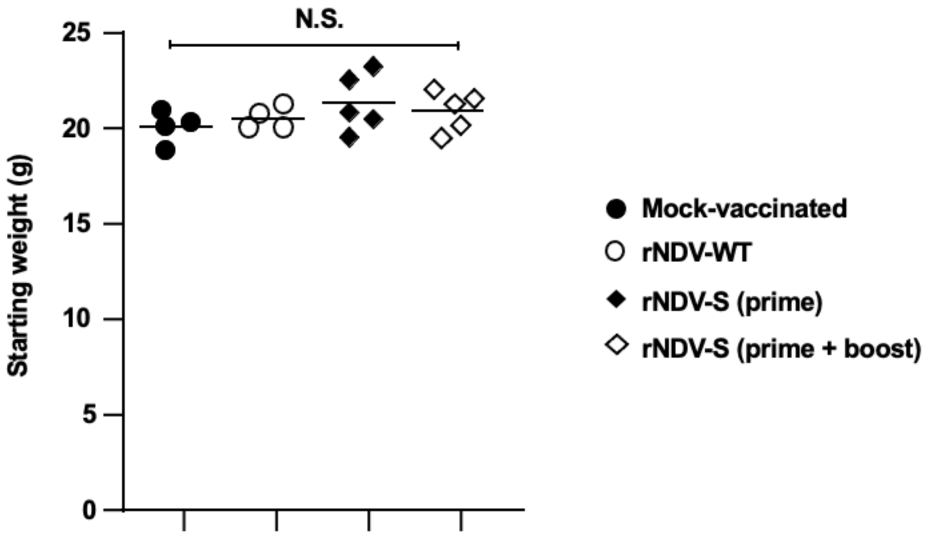


**Supplementary Figure 1. Starting weights did not significantly alter following intranasal rNDV-S vaccination.** Initial start weights of mice following instillation of PBS or indicated rNDV constructs intranasally over the experimental time-course. Data (n=4–5 mice/group); *, P<0.05; **, P<0.01; or ***, P<0.005 between naïve and infected groups, error bars represent SE of means via ANOVA with Dunnett's post-test.

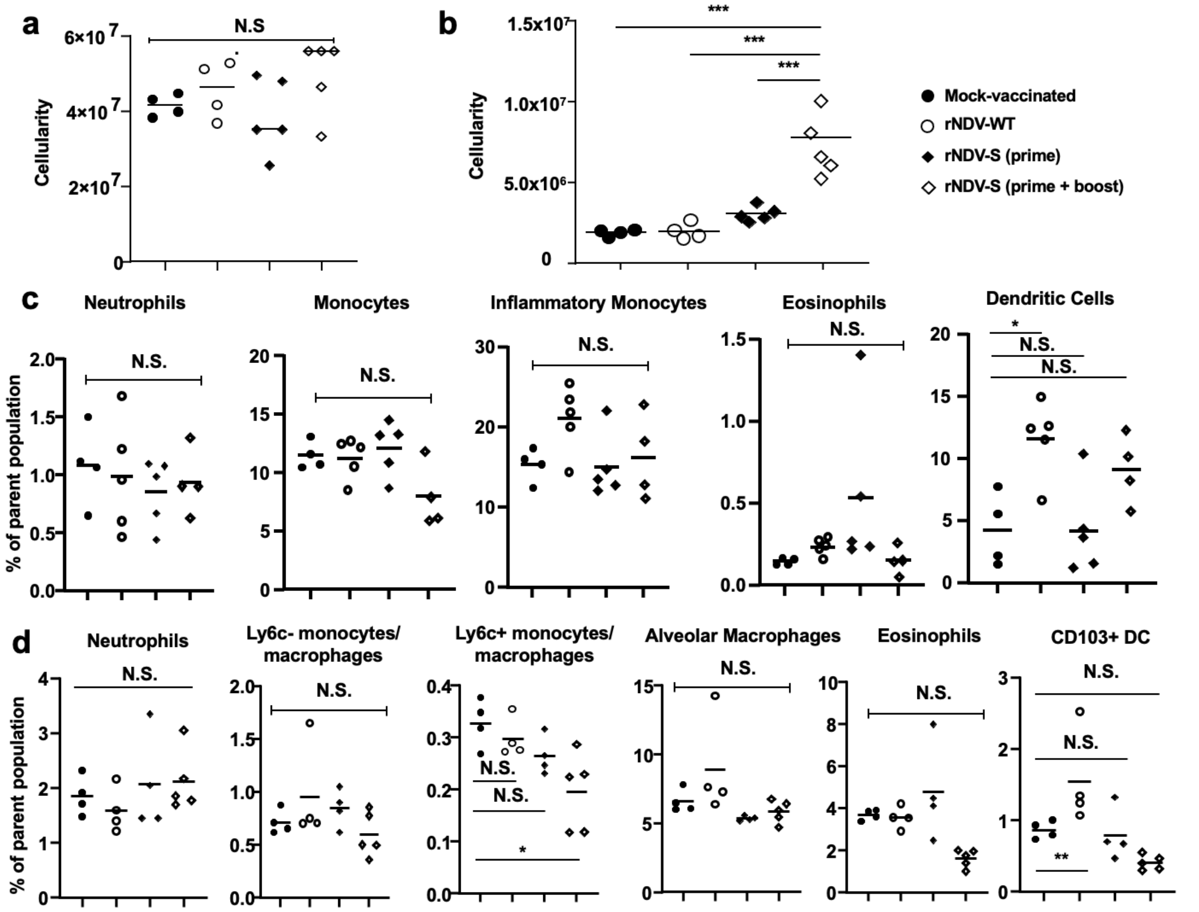


**Supplementary Figure 2. No adverse myeloid inflammatory response within the spleen or lung following vaccination with rNDV-S.** Cellularity of spleen (a), lung (b) and flow cytometric analysis of splenic (c) and lung (d) myeloid cell populations from mice on day 19 following instillation of mock PBS or indicated rNDV constructs intranasally on days 0 & 7. Data (n=4–5 mice/group); *, $P<0.05$; **, $P<0.01$; or ***, $P<0.005$ between naïve and infected groups, error bars represent SE of means via ANOVA with Dunnett's post-test.

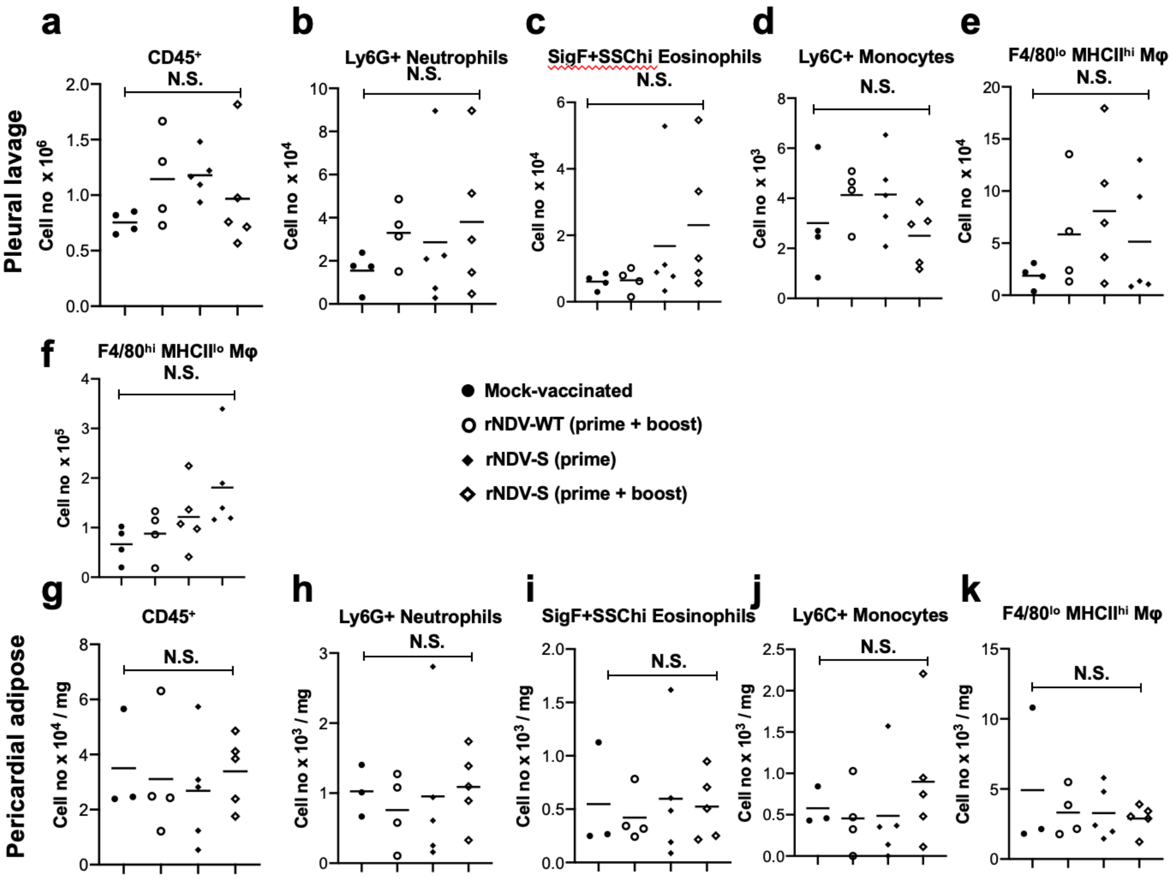


**Supplementary Figure 3. No lasting inflammatory response within the pleural and pericardial cavities following vaccination with rNDV-S:** Flow cytometric analysis of pleural lavage (upper panels) and digested total pericardial adipose (lower panels) cells from mice on day 19 following instillation of PBS or indicated rNDV constructs, on day 0 & 7. In Pleural lavage, total number of, CD45$^{+}$ cells **a)** Ly6G$^{+}$ Neutrophils (**b**) SigF$^{+}$SSC$^{hi}$ Eosinophils (**c**), Ly6C+ monocytes (**d**), F480$^{lo}$MHC-II$^{hi}$ Macrophages (**e**) and F4/80$^{hi}$MHC-II$^{lo}$ macrophages (**f**). In pericardial adipose, total number of, CD45$^{+}$ cells **g)** Ly6G$^{+}$ Neutrophils (**h**) SigF$^{+}$SSC$^{hi}$ Eosinophils (**i**), Ly6C+ monocytes (**j**) and F480$^{lo}$MHC-II$^{hi}$ Macrophages (**k**)

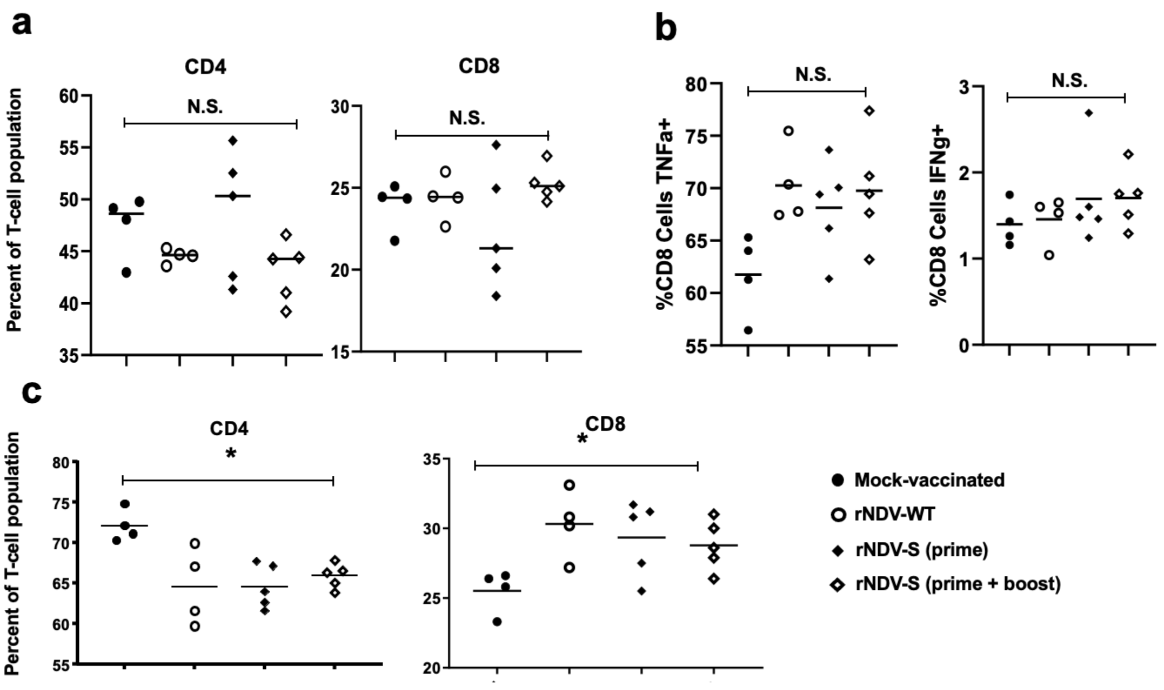


**Supplementary Figure 4. T-cell population dynamics and splenic T-cell cytokine production following vaccination:** Flow cytometric analysis of splenic T-cell populations (a), cytokine responses to SARS-CoV-2 spike protein (b) and (c) lung T-cell populations from mice on day 19 following instillation of mock PBS or indicated rNDVs constructs intranasally on days 0 & 7. Data (n=4–5 mice/group); *, P<0.05; **, P<0.01; or ***, P<0.005 between naïve and infected groups, error bars represent SE of means via ANOVA with Dunnett's post-test.

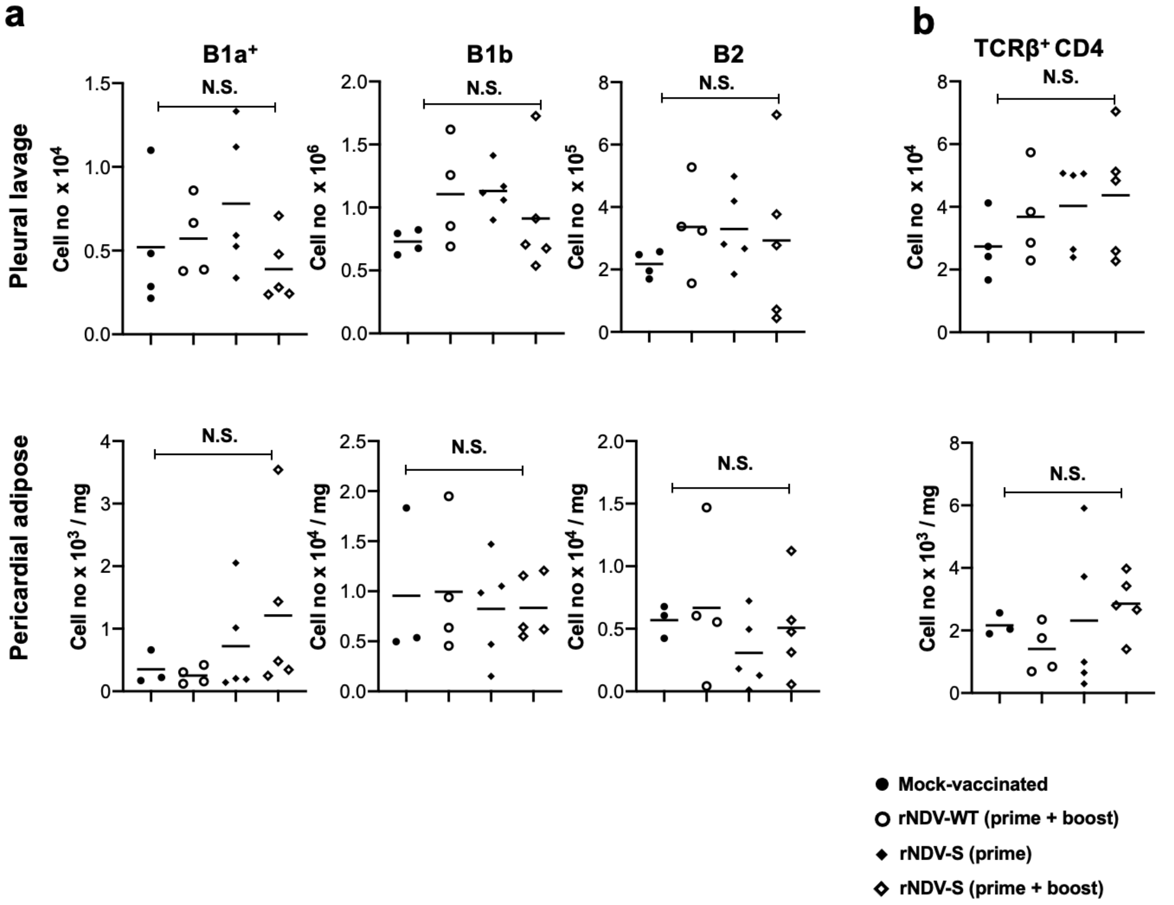


**Supplementary Figure 5**. **Adaptive immune cell populations within the pleural lavage and pericardial adipose a.** Total $CD19^+MHCII^+CD11b^+CD5^+$ B1a, $CD19^+MHCII^+CD11b^+CD5^-$ B1b & $CD19^+MHCII^+CD11b^-$ B2 cells, **b.** TCR**β**$^+$CD4$^+$ T cells determined by flow cytometric analysis of pleural lavage (upper panels) and digested total pericardial adipose (lower panels) cells from mice on day 19 following instillation of PBS or indicated rNDV constructs intranasally on days 0 & 7.

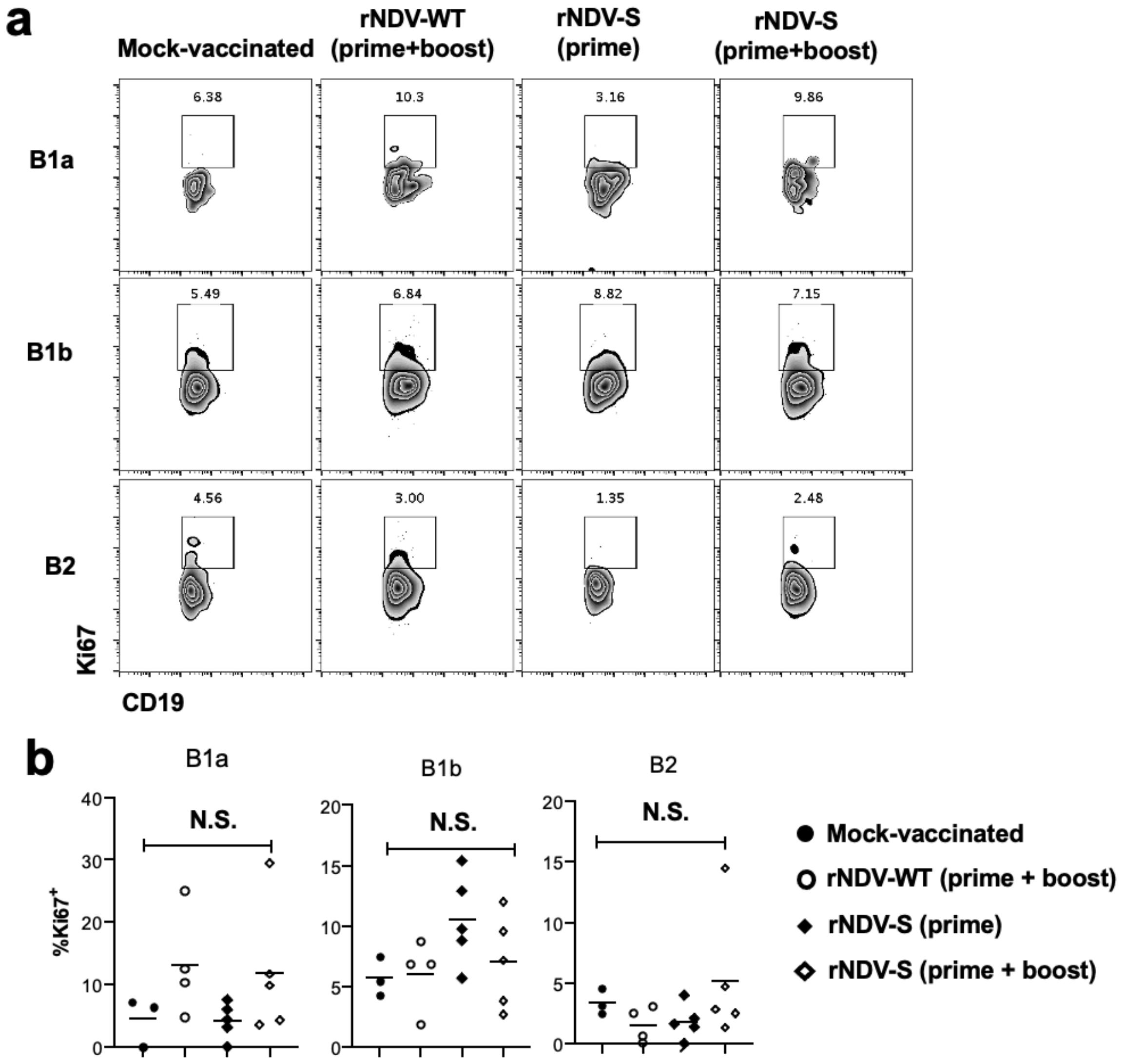


**Supplementary Figure 6. a**). Proliferation of pericardial B-cells isolated following digestion as determined via flow cytometric analysis on day 19 following instillation of PBS or indicated rNDV constructs intranasally on days 0 & 7. **b**) Quantitative presentation of different cell population shown in panel **a**.

### Prime groups

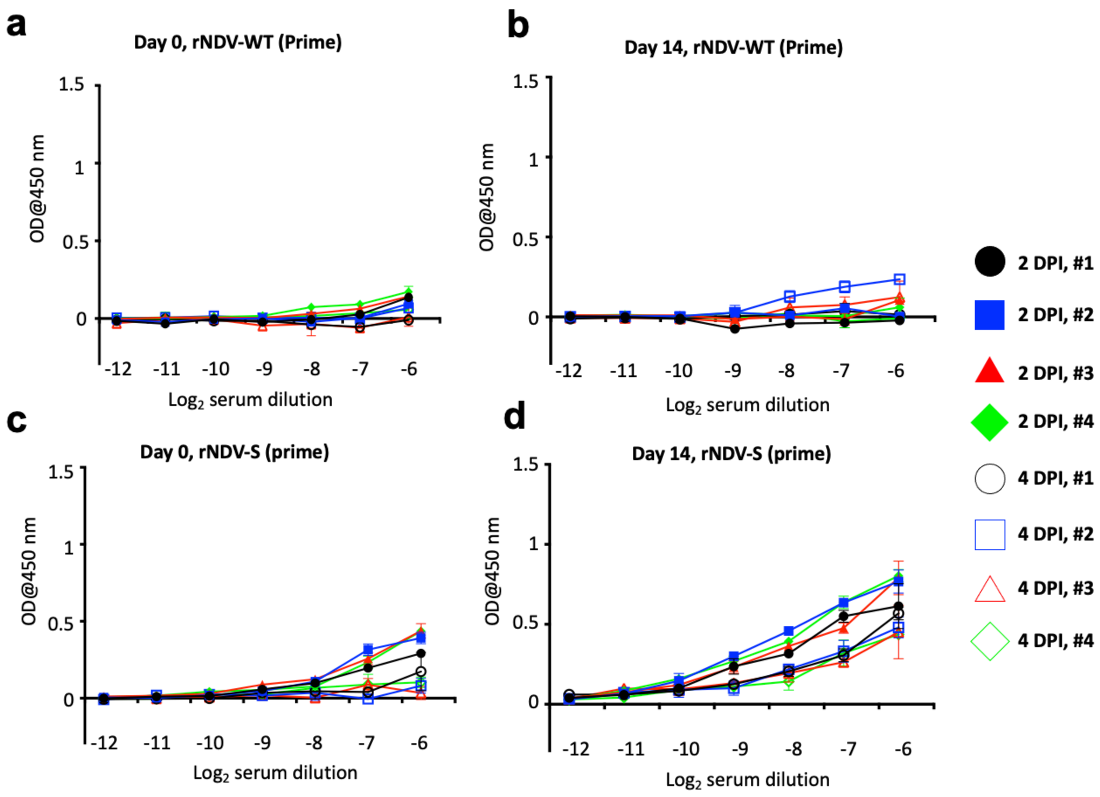


**Supplementary Figure 7. Total Abs in serum of individual golden Syrian hamsters vaccinated with rNDV constructs.** Levels of total S binding Abs in sera from individual hamsters vaccinated with either rNDV-WT or rNDV-S at 0 DPV (prime). Sera were collected at 0 (**a**) and 14 DPV (**b**) and individual hamster titre of Abs are displayed after 2 DPI and 4 DPI. The average of Abs titre in each group were presented in the Figure 6 in the main manuscript.

### Prime + boost groups

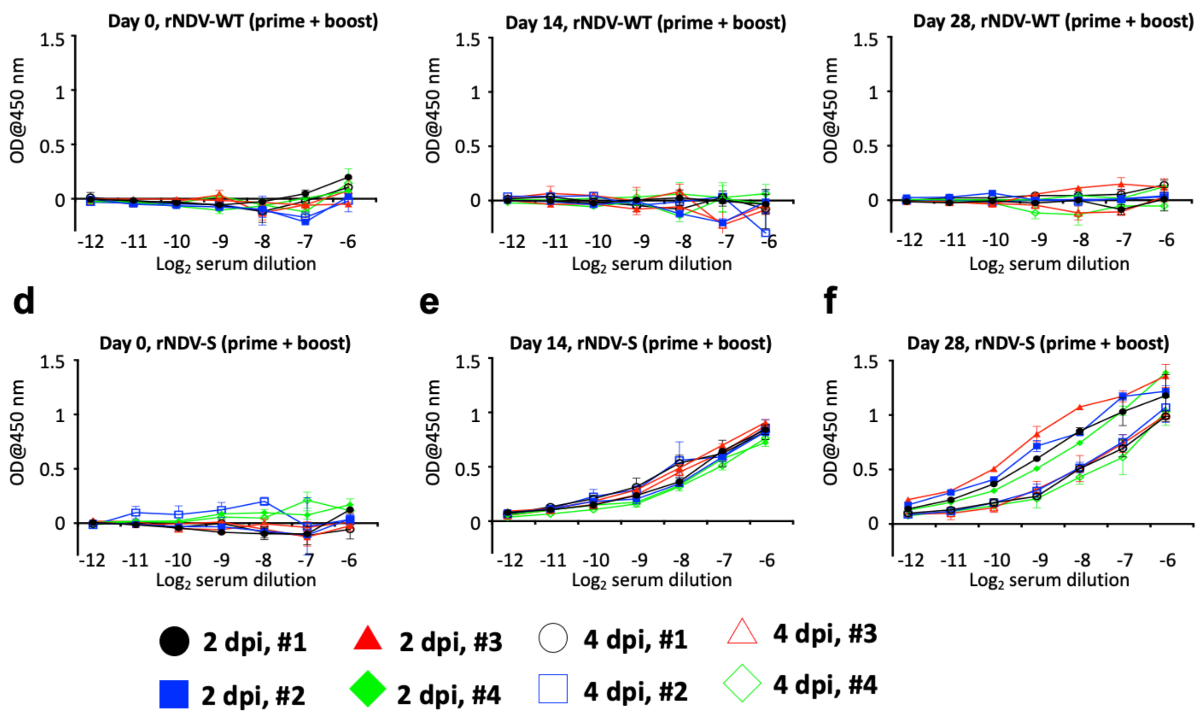


**Supplementary Figure 8. Total Abs in serum of individual golden Syrian hamsters vaccinated with rNDV constructs.** Levels of total S binding Abs in sera from individual hamsters vaccinated with either rNDV-WT or rNDV-S at 0 and 14 DPV (prime+boost). Sera were collected at 0 (**a**), 14 (**b**), and 28 DPV (**c**) and individual hamster titre of Abs are displayed after 2 DPI and 4 DPI. The average of Abs titre in each group were presented in the Figure 6 in the main manuscript.
